# Atg4 family proteins drive phagophore growth independently of the LC3/GABARAP lipidation system

**DOI:** 10.1101/2020.12.17.422596

**Authors:** Thanh Ngoc Nguyen, Benjamin Scott Padman, Susanne Zellner, Louise Uoselis, Marvin Skulsuppaisarn, Christian Behrends, Michael Lazarou

## Abstract

The sequestration of damaged mitochondria within double-membrane structures termed autophagosomes is a key step of PINK1/Parkin mitophagy. The Atg4 family of proteases are thought to regulate autophagosome formation exclusively by processing the ubiquitin-like Atg8 family (LC3/GABARAPs). We make the unexpected discovery that human Atg4s can directly promote autophagosome formation independently of their protease activity and of Atg8 family processing. High resolution structures of phagophores generated with artificial intelligence-directed 3D electron microscopy reveal a role for the Atg4 family in promoting phagophore-ER contacts during the lipid-transfer phase of autophagosome formation. Atg4 proximity interaction networks stimulated by PINK1/Parkin mitophagy are consistent with roles for Atg4s in protein/vesicle transport and lipid modification. We also show that Atg8 removal during autophagosome maturation does not depend on Atg4 de-lipidation activity as previously thought. Instead, we find that Atg4s can disassemble Atg8-protein conjugates, revealing a role for Atg4s as deubiquitinating-like enzymes. These findings establish non-canonical roles of the Atg4 family beyond the Atg8 lipidation axis and provide an AI driven framework for high-throughput 3D electron microscopy.

## INTRODUCTION

Macroautophagy (hereafter autophagy) is a lysosome-mediated degradative process critical for maintaining cellular homeostasis during nutrient deprivation and organelle damage. The Parkinson’s disease factors PINK1 and Parkin clear damaged mitochondria via a selective form of autophagy termed mitophagy (Lazarou et al., 2015; Narendra et al., 2008; Narendra et al., 2010). The kinase activity of PINK1 combines with the ubiquitin ligase Parkin to coat damaged mitochondria with ubiquitin chains (Kane et al., 2014; Koyano et al., 2014; Ordureau et al., 2014; Wauer et al., 2015). Ubiquitinated mitochondria are recognised by the autophagy receptors Optineurin (OPTN) and NDP52 which initiate phagophore formation by recruiting the ULK1 kinase complex (Lazarou et al., 2015; Vargas et al., 2019). It was recently shown that NDP52 recruits the ULK1 complex to mitochondria via its interaction with FIP200, this activates ULK1 kinase activity and initiates autophagosome biogenesis (Ravenhill et al., 2019; Vargas et al., 2019). ULK1 complex recruitment by NDP52 is facilitated by the kinase TBK1 which plays multiple roles in mitophagy including phosphorylation of autophagy receptors to promote ubiquitin binding (Heo et al., 2019; Moore and Holzbaur, 2016; Richter et al., 2016; Vargas et al., 2019).

The initiation of phagophore biogenesis and growth of phagophore membranes occurs on an ER derived membrane cradle termed the omegasome (Axe et al., 2008; Dikic and Elazar, 2018). Omegasomes are rich in the lipid PI3P which is generated by the class III phosphatidylinositol 3-kinase (PI3KC3) complex during nucleation of the phagophore membrane. Effector proteins including WIPI family members and DFCP1 bind to the omegasome via PI3P. WIPI1 and WIPI4 play a role in recruiting the lipid transfer proteins Atg2A and Atg2B which promote lipid transfer from the ER to the phagophore via contact sites (Kotani et al., 2018; Maeda et al., 2019; Osawa et al., 2019; Valverde et al., 2019; Zheng et al., 2017). The transmembrane protein Atg9A also functions to grow phagophore membranes by supplying proteins and lipid including the lipid kinase PI4kIIIb (Judith et al., 2019) at phagophore-ER contacts (Gomez-Sanchez et al., 2018). To date, high resolution quantitative measurements of phagophore-ER contacts during nucleation have been limited to 2D electron microscopy (EM) analyses (Gomez-Sanchez et al., 2018). Thus, phagophore-ER contacts have primarily been assessed by measuring a fraction of the phagophore and its surrounding environment. Current 3D analysis approaches using EM have been hindered by low throughput and difficulty of identifying nucleating phagophores. This has prevented quantitative analysis of phagophore-ER contacts which requires measurement of multiple phagophores from multiple cells.

Much still remains to be discovered about how phagophores grow to form autophagosomes, although the Atg8 machineries are thought to play key roles in membrane expansion, autophagosome-lysosome fusion, and degradation of the inner autophagosomal membrane (Abeliovich et al., 2000; Dikic and Elazar, 2018; Nakatogawa et al., 2007; Nguyen et al., 2016; Vaites et al., 2018). During PINK1/Parkin mitophagy, Atg8s drive ubiquitin independent recruitment of OPTN and NDP52 to phagophores to promote rapid autophagosome formation which amplifies mitophagy (Padman et al., 2019). There are six Atg8 family members in humans divided equally into the LC3 and GABARAP subfamilies consisting of LC3A, LC3B, LC3C and GABARAP, GABARAPL1 and GABARAPL2 respectively. Atg8s are attached to autophagosomal membranes via conjugation to the lipid phosphatidyl ethanolamine (PE) by the E3-like Atg12-Atg5-Atg16L1 complex (Ichimura et al., 2000; Mizushima et al., 1998). The conjugation machinery is recruited to phagophores by WIPI2b and the lipid PI3P (Dooley et al., 2014; Dudley et al., 2019). Atg8 lipidation is regulated by the Atg4 family of cysteine proteases which cleave Atg8s to expose a C-terminal glycine residue (Kirisako et al., 2000). The Atg8 lipidation reaction can also be reversed by Atg4s via proteolytic de-lipidation (Kauffman et al., 2018; Kirisako et al., 2000).

The Atg4 family consists of four members, Atg4A, Atg4B, Atg4C and Atg4B. Most of our understanding of Atg4 family function is centered on their protease activity toward Atg8 family members (Agrotis et al., 2019; Kauffman et al., 2018; Kirisako et al., 2000), which is regulated by post-translational modifications (Dikic and Elazar, 2018; Pengo et al., 2017; Sanchez-Wandelmer et al., 2017). In yeast, Atg4 mediated de-lipidation of Atg8 has been proposed to be critical for efficient autophagy by recycling Atg8s from mature autophagosomes and preventing non-specific lipidation of Atg8s on cellular membranes (Nair et al., 2012; Nakatogawa et al., 2012; Yu et al., 2012). The role of de-lipidation in human cells is less clear, although it has been shown that Ulk1 can phosphorylate Atg4B to prevent its proteolytic activity at phagophores (Pengo et al., 2017). This phosphorylation is important since excess Atg4 activity has been shown to negatively affect autophagy (Fujita et al., 2008a). The central role that Atg4s play in regulating autophagy coupled with their role in cancer have made their proteolytic activity an attractive drug target(Fu et al., 2019a; Fu et al., 2019b; Liu et al., 2018). A number of studies have assessed the *in vitro* proteolytic activity of Atg4s toward individual Atg8 family members helping to build a specificity profile of each Atg4 (Kauffman et al., 2018; Li et al., 2011). However, less is known about Atg4 activity in living human cells and this is largely due to the complexity of studying two protein families with multiple members. In addition, given that cellular stimuli can influence the activity of Atg4s (Betin and Lane, 2009; Perez-Perez et al., 2014; Scherz-Shouval et al., 2007), it is important to study the Atg4-Atg8 nexus in living cells under autophagy inducing conditions. It also remains to be determined whether Atg4 family specificity is driven by autophagy type, for example, PINK1/Parkin mitophagy.

In this study we discover a non-canonical role for the Atg4 family in phagophore growth during PINK1/Parkin mitophagy. Using multi-gene knockouts of the Atg4 and Atg8 families, we show that Atg4s can promote phagophore growth independently of their protease activity and of Atg8s by promoting phagophore-ER contacts. The proximity interaction proteome of Atg4s reveals a number of Atg4 associated processes including transport and lipid modification. We also show that Atg4s are dispensable for Atg8 de-lipidation from mature autophagosomes but are instead required to disassemble Atg8s conjugated to proteins in a post-translational modification we have termed Atg8ylation.

## RESULTS

### Specific Atg4 family members drive PINK1/Parkin mitophagy

To dissect the role of human Atg4 family members during PINK1/Parkin mitophagy, the genes encoding the four known Atg4s (Atg4A, Atg4B, Atg4C and Atg4D) were disrupted in HeLa cells using CRISPR/Cas9 gene editing (designated Atg4 tetra KO). Gene disruption was confirmed by sequencing of DNA indels (Table S1) and immunoblotting of Atg4A, Atg4B and Atg4C (Figure 1A). PINK1/Parkin mitophagy was assessed in Atg4 tetra KO cells by measuring the degradation of cytochrome *c* oxidase subunit II (CoxII), a mtDNA-encoded protein located within the mitochondrial inner membrane. To induce PINK1/Parkin mitophagy, mitochondria were damaged with Oligomycin and Antimycin A (OA). CoxII was robustly degraded in WT cells but not in Atg4 tetra KO cells (Figures 1B and 1C), which was consistent with the mitophagy block observed in cells lacking the Atg8 family (designated Atg8 hexa KO), and cells lacking Atg8 lipidation (Atg5 KO) (Figures 1B and 1C). Turnover of the mitochondrial outer membrane protein Mfn1 in all cell lines confirmed that Parkin activity was not disrupted (Figure 1B). To determine which individual Atg4 family members can drive PINK1/Parkin mitophagy, Atg4 tetra KO cells expressing YFP-Parkin were rescued with each mCherry tagged Atg4 family member (Figures 1D and 1E (see Figure 1G for mCherry-Atg4 expression levels)). Using the CoxII degradation assay, Atg4A, Atg4B and Atg4D restored mitophagy, whereas Atg4C did not (Figures 1D and 1E). To determine the mitophagy efficiency of each individual Atg4 protein, we also assessed mitophagy in Atg4 tetra KO rescue cell lines expressing YFP-Parkin and iRFP670 (hereafter iRFP) tagged Atg4s using the sensitive mtKeima mitophagy assay (Lazarou et al., 2015; Padman et al., 2019; Katayama et al., 2011). Atg4B was found to be less efficient than Atg4A and Atg4D in driving PINK1/Parkin mitophagy, whereas Atg4C showed little to no activity (Figure 1F). Taken together, these results define the Atg4 specificity and activity profile during PINK1/Parkin mitophagy and confirm the critical role that the Atg4-Atg8 axis plays in autophagy.

**Figure 1.**
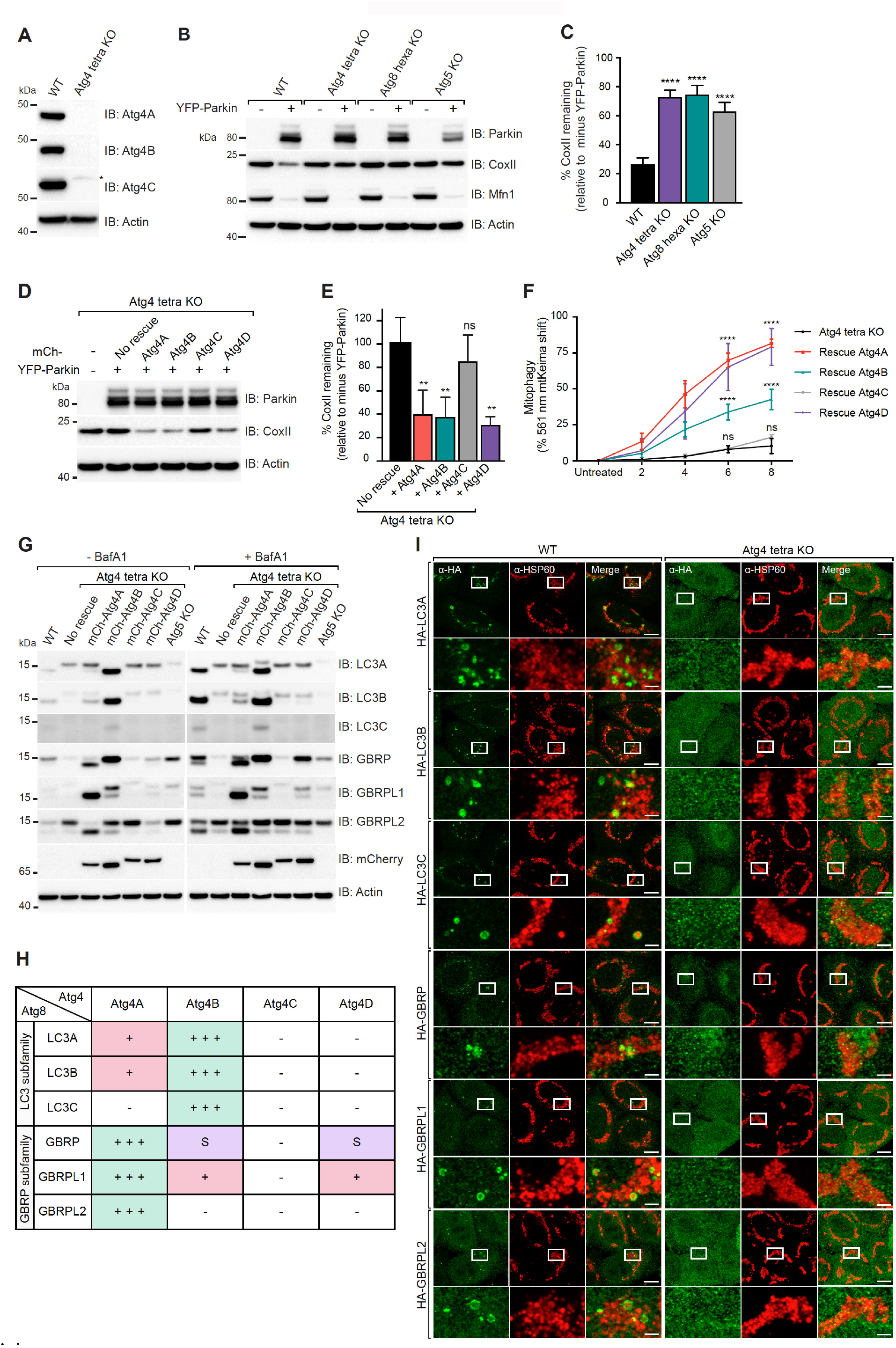
Atg4s are required for Pink1/Parkin mitophagy. (A) WT and Atg4 tetra KO were confirmed by immunoblotting (IB). kDa: kilodaltons. (B and C) WT, Atg4 tetra KO, Atg8 hexa KO and Atg5 KO with and without YFP-Parkin were treated with OA for 21 h and analysed by immunoblotting (B) and CoxII levels were quantified (C). (D and E) Atg4 tetra KO cells expressing YFP-Parkin and individual mCherry(mCh)-Atg4s were analysed by IB after 21 h OA treatment (D) and CoxII levels were quantified (E). (F) Atg4 tetra KO cells expressing YFP-Parkin, mtKeima and individual iRFP-Atg4s untreated or treated with OA for indicated times were analysed for lysosomal-positive mtKeima by FACS as the percentage of 561 nm mtKeima positive cells. (Representative FACS blots are provided in Figure S1B). (G and H) Atg4 tetra KO expressing YFP-Parkin without or with individual mCherry(mCh)-Atg4s were left untreated or treated with Bafilomycin A1 (BafA1) for 8 h. Cleavage of LC3s and GBRPs were then evaluated by immunoblotting (G) and summarized in (H). (-) indicates little to no cleavage activity whereas (+) in red rectangle and (+++) in green rectangle indicate weak and strong cleavage activity respectively. (S) in purple rectangle for stabilising effect on the substrate. (I) Representative images of WT and Atg4 tetra KO cells expressing YFP–Parkin and individual HA-LC3s or HA-GBRPs immunostained for HA and HSP60 after 3 h OA treatment. (untreated images shown in Figure S1A). Data in (C, E and F) are mean ± s.d. from three independent experiments. \**P*<0.05, \*\**P*<0.005, \*\*\**P*<0.001, \*\*\*\**P*<0.0001 (C, E one-way analysis of variance (ANOVA); F two-way ANOVA). ns: not significant. Scale bars: overviews, 10 µm; insets, 2 µm.

GABARAPs are the primary drivers of PINK1/Parkin mitophagy(Nguyen et al., 2016; Vaites et al., 2018), we therefore assessed the GABARAP and LC3 subfamily processing selectivity of each Atg4 family member. The cleavage/lipidation status of all six Atg8 family members was assessed in the presence or absence of 8 h Bafilomycin A (BafA1) treatment, in either WT or Atg4 tetra KO with or without rescue with each mCherry tagged Atg4 family member (Figures 1G and 1H). The appearance of lipidated Atg8 was interpreted as successful processing by an Atg4 family member. Atg5 KO cells were included as non-lipidated Atg8 controls (Figure 1G). Atg4A showed a preference for GABARAP subfamily members whereas Atg4B had a preference toward LC3s (Figures 1G and 1H). Atg4D primarily processed GABARAPL1, and had a stabilising effect on GABARAP levels, as did Atg4B (Figures 1G and 1H). LC3C and GABARAPL2 were specifically processed by Atg4B and Atg4A respectively (Figures 1G and 1H). Consistent with its putative role as a de-lipidation specialist (Kauffman et al., 2018), Atg4C did not robustly process any Atg8 family member (Figures 1G and 1H).

We next asked whether unprocessed Atg8s are recruited to mitochondria in the absence of Atg4 processing. Following 3 h OA treatment, all six HA-tagged Atg8 family members translocated to mitochondria in WT cells, but not in Atg4 tetra KOs (Figure 1I). Since Atg8 processing by Atg4s is required for lipidation, this result demonstrates that Atg8 recruitment during mitophagy is tied to lipidation rather than LIR mediated recruitment by autophagy receptors (Dooley et al., 2014; Fujita et al., 2008b; Padman et al., 2019).

### Atg4s are not essential for Atg8 family de-lipidation

In addition to processing Atg8s for lipidation, Atg4s have also been proposed to de-lipidate Atg8s to liberate them from PE. In yeast, de-lipidation is thought to contribute to autophagy by recycling Atg8s from mature autophagosomal structures (Nair et al., 2012; Nakatogawa et al., 2012; Yu et al., 2012). However, the role of de-lipidation in human cells is unclear. The role of de-lipidation was assessed by expressing mutants of Atg8 that bypass the need for Atg4 processing. These mutants have their C-terminal glycine residue exposed (Atg8-G) to enable lipidation in the absence of Atg4s. We confirmed that mutation of the C-terminal glycine does not affect the mitophagy activity of GABARAPs as assessed by CoxII degradation in Atg8 hexa KO cells (Figures S2E and S2F).

To explore the biological role of Atg4 mediated de-lipidation, we assessed the localisation of all six HA-Atg8-G family members under basal and mitophagy inducing conditions in the presence or absence of Atg4s. The analysis was conducted in WT, Atg4 tetra KO and Atg8 hexa KO cells, and a newly generated Atg4/Atg8 deca KO cell line (hereafter deca KO). The deca KO line (Figure 2A), was generated to eliminate the possibility that endogenous, unprocessed Atg8s have an influence on HA-Atg8-G structures. In untreated cells lacking the Atg4 family (Atg4 tetra KOs and deca KOs), HA-LC3A-G, HA-LC3B-G, HA-LC3C-G and HA-GABRAPL2-G were localised diffusely throughout the cytosol (Figures S2B and S2D), whereas HA-GABARAP-G and HA-GABARAPL1-G were additionally localised to the Golgi compartment (Figures S2B, S2D and S2H). Golgi localisation of HA-GABARAP-G and HA-GABARAPL1-G was not observed in WT or Atg8 hexa KO cells which express endogenous Atg4s (Figures S2A and S2C). Thus, Atg4s basally prevent non-specific accumulation of GABARAP and GABARAPL1 on the Golgi. Upon mitophagy induction (3 h OA treatment), all six Atg8-Gs were recruited to mitochondria in deca KO cells (Figure 2B), and in Atg4 tetra KO cells (Figure S2B), although there appeared to be fewer Atg8-G structures on mitochondria in comparison to WT and Atg8 hexa KO (Figures S2A and S2C). This result indicates a role for Atg4 de-lipidation activity in mitophagy. We therefore assessed the PINK1/Parkin mitophagy activity of HA-GABARAPL1-G in the presence and absence of Atg4s using the mtKeima mitophagy assay. HA-GABARAPL1-G was chosen because it is highly active during PINK1/Parkin mitophagy(Nguyen et al., 2016). In the presence of Atg4s (Atg8 hexa KOs), the HA-GABARAPL1-G bypass mutant had robust levels of mitophagy, which were similar to WT GABARAPL1 (Figure 2C). In contrast, mitophagy was significantly lower in deca KOs lacking Atg4s (Figure 2C). Taken together, these results indicate that Atg4 activity is crucial for efficient mitophagy.

**Figure 2.**
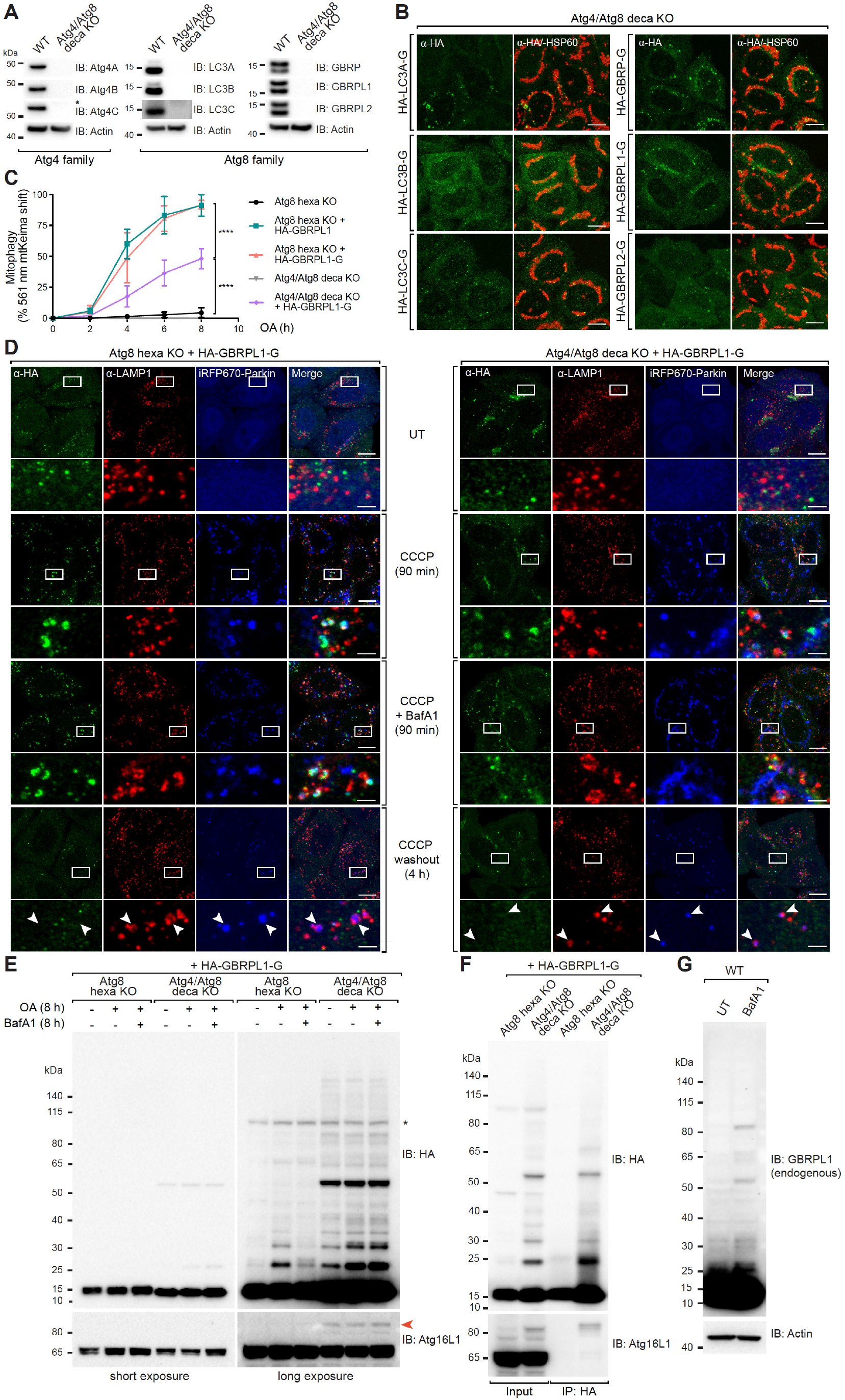
Atg4s are not essential for de-lipidation, but can reverse conjugation of Atg8s to cellular proteins. (A) Atg4/Atg8 deca KO were confirmed by immunoblotting (IB). kDa: kilodaltons. (B) Representative images of Atg4/Atg8 deca KO cells expressing iRFP–Parkin and individual HA-tagged Atg8-G mutants that do not require Atg4 initial cleavage, immunostained for HA and HSP60 after 3 h OA treatment. (untreated images shown in Figure S2D). (C) FACS analysis of the percentage of 561 nm mtKeima-positive cells for Atg8 hexa KO cells and Atg4/Atg8 deca KO cells expressing mtKeima, BFP-Parkin and indicated form of GBRPL1 following time course incubation with OA. (Representative FACS blots are provided in Figure S2G). (D) Atg8 hexa KO and Atg4/Atg8 deca KO cells expressing iRFP-Parkin and HA-GBRPL1-G were incubated with DMSO or 10 μm CCCP in the absence or presence of BafA1 for 90 min. After CCCP incubation, CCCP was washed out and cells were left growing in full medium for 4 h. Cells were immunostained for HA and mitochondrial HSP60 before analysed by confocal imaging. (E) Cell lysates from Atg8 hexa KO and Atg4/Atg8 deca KO cells expressing iRFP-Parkin and HA-GBRPL1-G untreated or treated with OA for 8 h in the absence or presence of BafA1 were analysed by immunoblotting for HA and Atg16L1. Red arrow and asterisk denote Atg16L1-GABARAPL1 conjugate and non-specific band respectively. (F) Western blot analysis of denaturing HA immunoprecipitation performed on untreated lysates from Atg8 hexa KO and Atg4/Atg8 deca KO cells expressing iRFP-Parkin and HA-GBRPL1-G. (G) Lysate samples from WT untreated and treated with BafA1 were immunoblotted for endogenous GBRPL1. Data in (C) are mean ± s.d. from three independent experiments. \*\*\*\**P*<0.0001 (two-way ANOVA). Scale bars: overviews, 10 µm; insets, 2 µm.

We hypothesised that Atg4 activity regulates mitophagy by either de-lipidating Atg8s, or by playing a direct role in autophagosome formation. Given that Atg8 de-lipidation from mature autophagosomal structures is important for autophagy in yeast (Nair et al., 2012; Yu et al., 2012), we first assessed whether GABARAPL1-G becomes trapped on mature autophagosomes in the absence of Atg4s. Following mitophagy activation with CCCP, HA-GABARAPL1-G localised to iRFP-Parkin coated mitochondria, some of which were positive for LAMP1, in both Atg8 hexa KOs and deca KOs (Figure 2D, CCCP (90 min) panel). Simultaneous treatment with CCCP and BafA1 resulted in an accumulation of mitochondrial HA-GABARAPL1-G structures including autolysosomes (Figure 2D, CCCP + BafA1 (90 min) panel), confirming that HA-GABARAPL1-G labelled mitophagosomes undergo turnover by lysosomes.

Next, we assessed HA-GABARAPL1-G de-lipidation from mature autophagosomal structures including autolysosomes. Cells were treated with CCCP followed by a 4 h washout period to switch mitophagy off and enable turnover of autolysosomes. HA-GABARAPL1-G was absent from autolysosomes in Atg8 hexa KOs as evidenced by the lack of HA-GABARAPL1-G on iRFP-Parkin and LAMP1 double-positive structures (Figure 2D, CCCP washout panel, white arrows). In addition, few to no iRFP-Parkin labelled mitochondria had HA-GABARAPL1-G on their surface (Figure 2D, CCCP washout panel). In contrast to the expected role for Atg4s in de-lipidation, HA-GABARAPL1-G was removed from autolysosomes in deca KOs lacking Atg4s (Figure 2D, CCCP washout panel, white arrows). Thus HA-GABARAPL1-G can be removed from mature autophagosomes in the absence of Atg4s. Similar results were observed in starved cells that were then fed to induce Atg8 recycling (Figure S3A). Atg4 de-lipidation activity was also dispensable for mitophagosome formation (Figure S2I). Taken together, these results argue against a critical role for Atg4 mediated de-lipidation in autophagosome formation and Atg8 recycling in human cells.

### Deubiquitin-like activity of Atg4s

During our analyses of HA-GABARAPL1-G, HA-LC3B-G and HA-GABARAP-G, we observed the formation of high molecular weight species on SDS-PAGE which increased in intensity and number in deca KOs lacking Atg4s (Figures 2E and S3B). In addition, a higher molecular weight species of Atg16L1 was identified in deca KOs expressing HA-GABARAPL1-G (Figure 2E, red arrow), its size consistent with the attachment of a 15 kDa species of HA-GABARAPL1-G. The higher molecular weight species of HA-GABARAPL1-G and Atg16L1 were resistant to the deubiquitinating enzyme USP2 (Figure S3C), eliminating the possibility that they are ubiquitinated forms of the proteins. To confirm that HA-GABARAPL1 is covalently attached to proteins including Atg16L1, a denaturing co-immunoprecipitation of HA-GABARAPL1-G was performed. The co-immunoprecipitation revealed that the high molecular weight species of Atg16L1, which accumulated in the absence of Atg4s, is a covalent conjugate of HA-GABARAPL1-G (Figure 2F). HA-GABARAPL1-G was also covalently attached to a number of unidentified proteins, as evidenced by the high molecular weight smears in deca KOs (Figures 2E and 2F). Re-introduction of Atg4A or Atg4B, but not Atg4C or Atg4D, reduced the high molecular weight smears of HA-GABARAPL1-G (Figure S3D). The lack of Atg4D activity in reducing HA-GABARAPL1 conjugates is in contrast to its activity toward cleaving GABARAPL1 (Figures 1G and 1H). The high molecular weight conjugates of HA-GABARAPL1-G are reminiscent of poly-ubiquitin smears, and given that Atg8s are ubiquitin-like proteins, we have termed the covalent attachment of Atg8s to proteins Atg8ylation. Atg8ylation is unlikely to be a spurious event resulting from ectopic expression of HA-GABARAPL1-G since high molecular weight forms of endogenous GABARAPL1 were also observed in BafA1 treated WT cells (Figure 2G). Taken together, our results show that although Atg4s are not critical for Atg8 de-lipidation during mitophagy, they can function as deubiquitinating-like enzymes to regulate the levels of Atg8ylation.

### Atg4s play a direct role in autophagosome formation independent of Atg8s

Despite being dispensable for Atg8 recycling (Figures 2D and S3A), Atg4s are required for efficient mitophagy (Figure 2C). We therefore investigated whether Atg4s play a direct role in autophagosome formation. EM analysis of Atg4 tetra KO cells revealed a significant mitophagosome formation defect compared to WT controls (Figures 3A and 3B). The Atg4 tetra KO mitophagosome defect is in conflict with previous studies showing that autophagosomal structures can form in cells lacking Atg8s (Nguyen et al., 2016; Vaites et al., 2018), or lacking Atg8 lipidation (Tsuboyama et al., 2016) (Figure S4A). A possible explanation for the mitophagosome defect in Atg4 tetra KOs is that Atg4s play an Atg8 independent role during autophagy. Alternatively, the presence of unprocessed Atg8s in Atg4 tetra KO cells may inhibit mitophagosome formation. To exclude the latter possibility, mitophagosome formation was compared between Atg8 hexa KOs and deca KOs. It is important to note that the Atg8 hexa KO line is the parental line from which the deca KO line was generated. As can be seen in Figures 3C and 3D, mitophagosomes form in the absence of Atg8s (Atg8 hexa KO), but do not form upon the deletion of Atg4s (deca KO). Consistently, fewer Atg13 labelled phagophores (Figures S4B and S4C), and WIPI2 labelled omegasomes (Figures 3E and 3F) were observed in deca KOs compared to Atg8 hexa KO controls. Reintroduction of all for Atg4s in deca KOs restores the number and size of WIPI2 foci (Figures 3E-3G), as does the reintroduction of catalytically inactive Atg4s. These results demonstrate that Atg4s can promote autophagosome biogenesis independently of their protease activity and of Atg8s.

**Figure 3.**
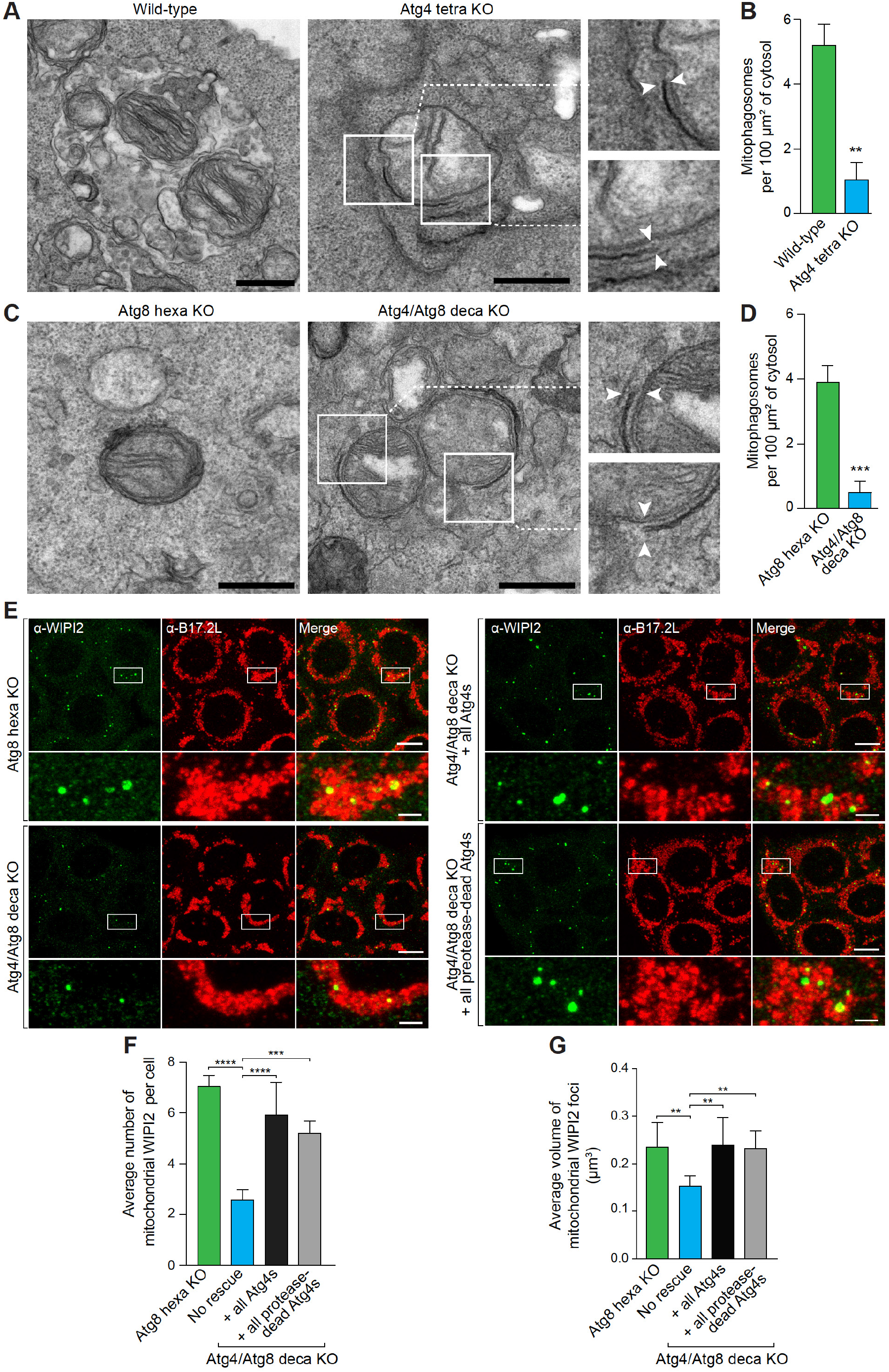
Atg4s promote autophagosome formation independent of their protease activity and of Atg8s. (A and B) WT and Atg4 tetra KO expressing YFP-Parkin were subjected to 6 h treatment with OA and BafA1 and subsequent TEM analysis (A) and the average number of mitophagosomes per 100 μm^2^ of cytosol was quantified (B). (C and D) Atg8 hexa KO and Atg4/Atg8 deca KO cells expressing YFP-Parkin were subjected to 6 h treatment with OA and BafA1 and subsequent TEM analysis (C) and the average number of mitophagosomes per 100 μm^2^ of cytosol was quantified (D). (E-G) Atg8 hexa KO, Atg4/Atg8 deca KO and Atg4/Atg8 deca KO cells rescued with all wild type Atg4s or all protease-dead Atg4s (all cell lines express iRFP-Parkin) were treated with OA for 3 h and immunostained for WIPI2 (E). The average number of WIPI2 mitochondrial structures per cell (F) and the average volume of WIPI2 mitochondrial foci (G) were analysed. Data in (B, D, F, G) are mean ± s.d. from three independent experiments; \*\**P*<0.005, \*\*\**P*<0.001, \*\*\*\**P*<0.0001 (B, D one-way ANOVA; F, G two-way ANOVA). Scale bars: (A and C), 500 nm; (E), overviews, 10 µm; insets, 2 µm.

### Atg4 proximity interaction networks reveal their activity during phagophore nucleation

To understand how Atg4s might contribute to autophagosome formation during mitophagy, we sought to determine which autophagosome intermediates the Atg4s associate with. Atg4 family members expressed in Atg4 tetra KOs did not translocate to mitochondria during mitophagy (Figure S5A). The addition of wortmannin, which accumulates early phagophore structures also did not stabilise Atg4s on mitochondria (Figure S5A). An APEX2 based proximity biotinylation system was used instead to enable detection of labile Atg4 interactions and to map the proximity interaction network of each Atg4. Individual Myc-APEX2-Atg4s were stably expressed in Atg4 tetra KO cells (Figure S5B). Myc-APEX2-Atg4A, -4B and -4D were functional in rescuing PINK1/Parkin mitophagy, whereas APEX2-Atg4C showed minimal rescue (Figure S5C), consistent with earlier observations (Figures 1D and 1E).

To determine the proximity interaction network of Atg4s during mitophagy, Myc-APEX2-Atg4 cell lines were either untreated, treated with OA, or treated with wortmannin and OA (Wort/OA). Biotinylated proteins were captured and subjected to mass spectrometry analysis. A robust increase in the number of biotinylated proteins was observed upon mitophagy activation for Atg4A, B and D, but not Atg4C (Figures 4A, 4B, S5D and S5E). Among the OA-induced proximity targets, Myc-APEX2-Atg4A, -Atg4B and -Atg4D uniquely biotinylated 124, 38 and 57 proteins respectively and shared 24 common targets (Figure 4B). There were also a number of targets that were commonly found in two of the three mitophagic Atg4s, with 74 targets shared by Atg4A and Atg4D, 34 shared by Atg4A and Atg4B and 11 shared by Atg4B and Atg4D (Figure 4B). This indicates that although Atg4s can redundantly contribute to autophagosome formation through common factors, they can also function non-redundantly through exclusive interaction networks. Functional annotation clustering of the combined Atg4 proximity network revealed pathways associated with autophagy, endosomal and vesicle transport, multivesicular body assembly and protein transport (Figure 4D).

**Figure 4.**
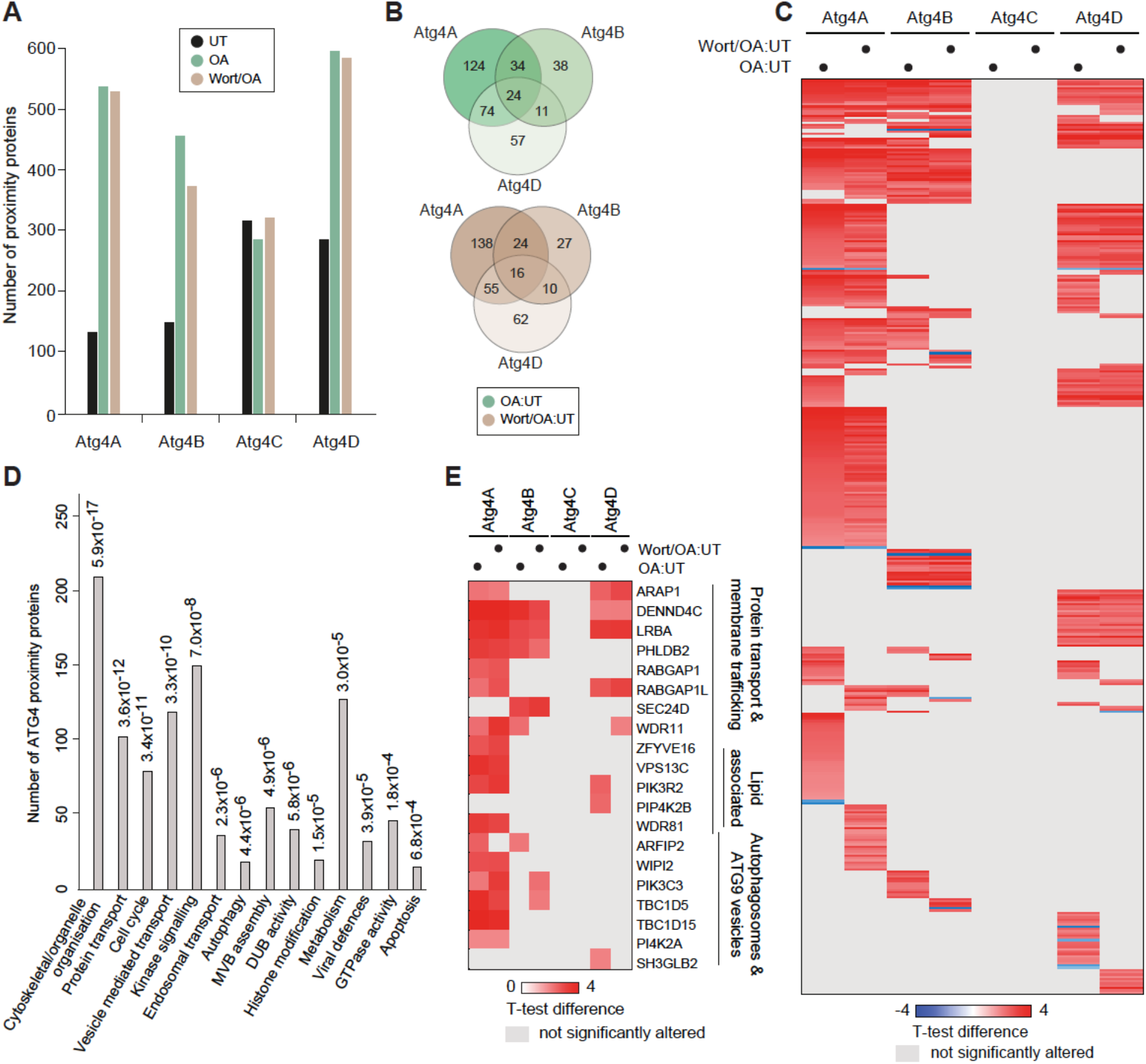
Proximity biotinylation reveals early autophagy machinery near APEX2-Atg4s during mitophagy induction. (A-E) Atg4 tetra KO cells stably expressing YFP-Parkin without or with individual APEX2-Atg4s were left untreated or treated with OA or Wort/OA for 3 h in triplicates. Following biotin labelling reactions, cells were harvested, subjected to lysis and streptavidin pulldown prior to MS analysis. (A) The bar graph shows the total number of proximity proteins which were identified in each condition (raw data shown in Table S4). (B) Overlap of proximity proteins that are significantly changed upon OA and Wort/OA treatment between Atg4A, Atg4B and Atg4D. (C) Heat map of proximity proteins whose abundance significantly change upon OA and Wort/OA. (D and E) Gene Ontology analysis (D, raw data shown in Table S5) and heat map representation of some representatives (E) of proteins from (C). For (E) and (C) student t-test was used to determine the statistical significance of the abundance alterations of biotinylated proteins between treatments (OA vs. untreated or OA/Wortmannin vs. untreated). A p-value ≤ 0.05 was considered statistically significant.

Comparing mitophagy induced Atg4 proximity networks in the presence or absence of wortmannin can provide information about Atg4 phagophore association before and during PI3P production. The addition of wortmannin during mitophagy induction abolished the significant enrichment of a number of proteins in the proximity of Atg4s (Figures 4C and 4E). For example, wortmannin treatment either prevented or decreased proximal interactions between both Atg4A and Atg4B with the Atg9 vesicle associated factor ARFIP2 (Judith et al., 2019), and Atg4A with the Atg9a vesicle transport factor TBC1D5 (Popovic and Dikic, 2014) (Figure 4E). In contrast, there were a subset of proteins whose proximity to Atg4s increased in the presence of wortmannin. These included the PI3P kinase PIK3C3 (Vps34) for Atg4A and Atg4B, TBC1D5 and SEC24D (Stadel et al., 2015) for Atg4B, WDR11 for Atg4A and Atg4D (Figure 4E), and FYCO1 (Pankiv et al., 2010) and NBR1 (Kirkin et al., 2009) for Atg4A (Table S4). Overall, the wortmannin analyses demonstrate that the majority of mitophagy induced Atg4 proximity networks form independently of PI3P production. This places Atg4 function at a very early stage of autophagosome formation during phagophore nucleation.

### Computer vision reveals that Atg4s promote phagophore growth and phagophore-ER contacts during the lipid transfer phase of autophagosome formation

The proximity interactome of Atg4s during mitophagy revealed factors associated with Atg9 vesicles (ARFIP2, PI4K2A, TBC1D5), phagophore nucleation including the omegasome ER cradle (WIPI2, PIK3C3, TBC1D15), trafficking and lipid modification (PIK3R2, PIP4K2B, INPP4A, PI4K2A) (Figure 4E and Table S4). These networks led us to hypothesise that Atg4s promote lipid transfer/transport to the growing phagophore during nucleation and therefore may play a role in phagophore-ER contacts. To explore this hypothesis, we utilised Focused Ion Beam-Scanning Electron Microscopy (FIB-SEM) to image phagophore ultrastructure and phagophore-ER contacts at high-resolution in 3D.

FIB-SEM is ideally suited for the acquisition of high-resolution 3D image data from large cellular volumes (Hoffman et al., 2020). However, traditional methods for the reconstruction and analysis of such data requires manual image segmentation and classification. The hand-drawn surfaces that are generated during manual segmentation are laborious to produce, low throughput and subject to human imprecision, making them unsuitable for large scale morphometric analyses. To overcome these limitations, we developed an approach which delegates the most time-consuming tasks to an artificial intelligence (AI) within a newly developed framework that captures the true surface detail of intracellular organelles. Using the machine learning capabilities of Waikato Environment for Knowledge Analysis (WEKA) in combination with ImageJ (Arganda-Carreras et al., 2017; Frank et al., 2004a; Schindelin et al., 2012), we trained a random forest computer vision model to faithfully detect all cellular membranes in volumetric FIB-SEM data (Figures 5A and 5B). However, the binary segmentation data from the AI was not directly used to triangulate 3D surfaces for membranes. Instead, we used the computer vision data to selectively extract membranes directly from the EM data (Figure 5B). By doing so, the intensity of membrane voxels from the EM data contributed directly to surface triangulation which generates 3D rendering of surfaces that faithfully capture the true texture and detail of membranes (Figure 5C). We have termed this new 3D framework AIVE (AI-directed Voxel Extraction).

**Figure 5.**
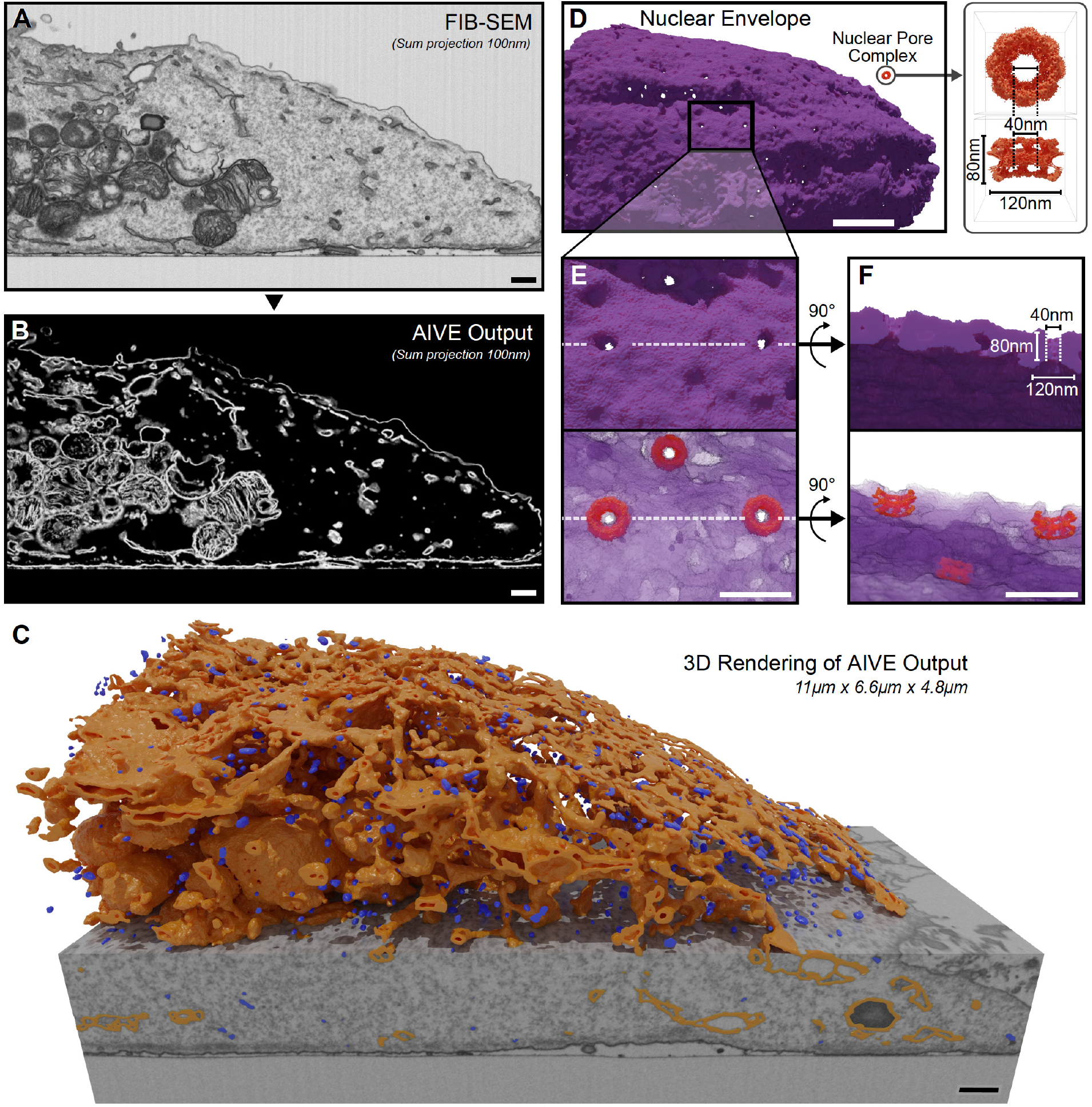
An AI-directed approach to the detection and rendering of membranes in volumetric EM data. (A and B) FIB-SEM data acquired from a HeLa cell (A), with representative outputs generated by AI-directed Voxel Extraction (AIVE; B). (C) 3D rendering of all membranes detected using AIVE (other than the plasma membrane), with vesicles (blue) separated from other membranes (gold) by size (C). (D) Sub-classified membranes of the nuclear envelope shown (to scale) alongside the Cryo-EM structure of the human Nuclear Pore Complex solved *in situ* by A von Appen et al., 2015. (E and F) 3D renderings of the nuclear pore complex (to scale) overlaid with the nuclear envelope, viewed from the front (E), or rendered with a cutaway from below (F). Scale bars: (A-D), 500 nm; (E and F), 200 nm.

To demonstrate the precision and accuracy of AIVE, we compared manually segmented surfaces of the nuclear membrane (Figure S7A), with AI segmentation alone (Figure S7B), and then with AIVE (Figure S7C). Note the lack of terracing artefacts and the substantial improvement in surface detail in Figure S7C. To further demonstrate the capabilities of our approach, the size of nuclear pores detected by AIVE were compared with the *in situ* cryo EM structure of the nuclear pore complex (von Appen et al., 2015) (Figure 5D). The dimensions of nuclear pores detected using AIVE agree with those of the nuclear pore complex (Figures 5E and 5F and Movie S1). These results demonstrate the precision and accuracy of our AI-directed approach, which is required for high-throughput quantitative assessments of phagophore morphology and contacts with the ER.

To identify phagophores for 3D reconstruction, we utilized correlative light electron microscopy (CLEM). The lipid transfer protein Atg2B was chosen as the correlative imaging target (Figures 6A and 6B) because it sits at the nexus of omegasome formation, lipid transfer and Atg9 vesicles. To focus on Atg8 independent functions of the Atg4s, phagophore structures were compared between the hexa KOs and deca KOs (Figures 6C and 6D). Reconstructions of GFP-Atg2B labelled phagophores in Atg8 hexa KOs (Figures 6E, 6F, S8A and S8B, and Movies S2 and S3) show mitochondria (red) in the process of phagophore engulfment (green). Phagophore contacts with ER membranes (blue) were also observed (Figures 6E, 6F, and Movies S2 and S3), consistent with the role of Atg2s in lipid transfer from ER membranes (Gomez-Sanchez et al., 2018; Kotani et al., 2018; Osawa et al., 2019; Valverde et al., 2019). However, in the absence of Atg4s (deca KOs), phagophores were smaller (Figures 6G-6I, S8C, and S8D), and had less ER occupying the space surrounding the phagophore (Figures 6G, 6H and 6J and Movies S4 and S5). Phagophores in deca KOs also had decreased surface roughness, i.e. fewer pits and protrusions (Figure 6K), which may correlate with the decrease in ER contacts. Consistent with the phagophore formation defect, fewer GFP-Atg2B structures were observed in confocal microscopy analyses of deca KOs compared to Atg8 hexa KOs (Figures S4D and S4E). These results show that Atg4s promote phagophore growth and phagophore-ER contacts during the Atg2B lipid transfer phase of autophagosome formation. Notably, this function of Atg4s is independent of Atg8s.

**Figure 6.**
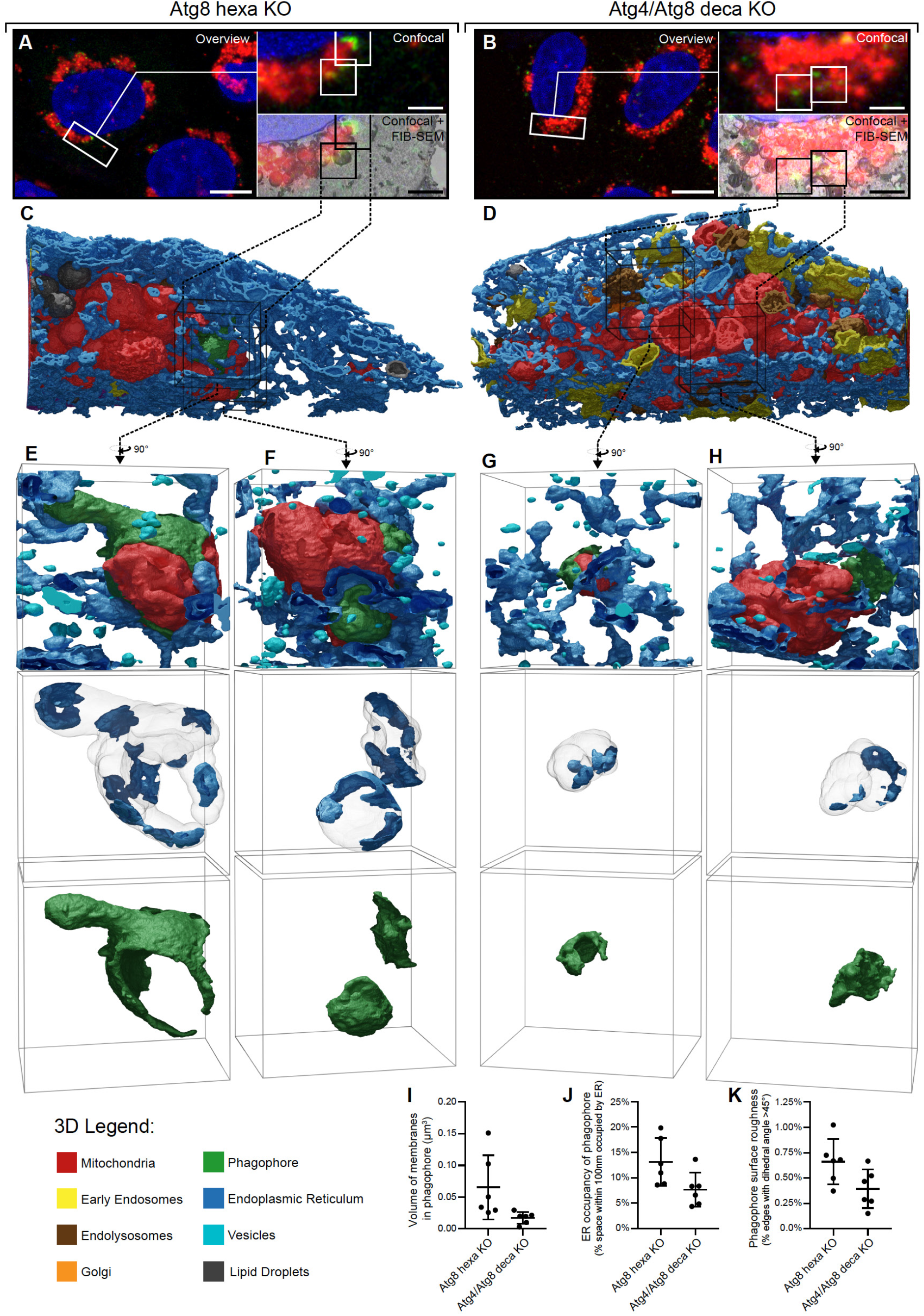
Atg4s directly promote phagophore growth and phagophore-ER contacts during PINK1/Parkin mitophagy. (A and B) Correlative alignment between the deconvolved optical data and orthosliced (X-Z) FIB-SEM data, showing the regions of interest in Atg8 hexa KO (A) and Atg4/Atg8 deca KO (B) cells expressing Mito-DsRed, iRFP-Parkin and GFP-Atg2B after 3 h incubation with OA. (C and D) 3D rendering of all membranes detected within the volume-of-interest in Atg8 hexa KO (C) and Atg4/Atg8 deca KO (D) with insets (white boxes) indicating the position of Atg2B positive phagophores. (E-H) 3D renderings of insets displaying neighbouring endomembranes (upper insets), the ER detected within 100 nm of the phagophores (middle insets), and Atg2B positive phagophores (lower insets) from the Atg8 hexa KO (E and F) and Atg4/Atg8 deca KO (G and H) cell lines (animated renderings of insets provided in Movies S2-S5). (I-K), 3D morphometric surface analysis of the Atg2B positive phagophores quantifying the volume of phagophore membrane (I), the proportion of space near the phagophore (<100 nm) occupied by ER (J), and the local roughness of the phagophores surfaces (K). Data in (I) and (J) are raw measurements with mean ± s.d. for 6 phagophores (two cells per condition, 3 confirmed Atg2B positive phagophores per cell; All phagophores shown in Figure S8). *P<0.05 (students t-test). Scale bars: optical overviews, 10 µm; insets, 2 µm; inset cubes, 2 µm x 2 µm x 2 µm.

## DISCUSSION

The Atg8 conjugation system has long been recognised as playing central roles in all forms of canonical autophagy (Dikic and Elazar, 2018; Mizushima et al., 1998). In human cells, Atg8 conjugation to PE promotes the expansion of autophagosomal membranes, autophagosome-lysosome fusion, and degradation of the inner autophagosomal membrane (Nguyen et al., 2016; Tsuboyama et al., 2016; Vaites et al., 2018). Human Atg4s are tasked with governing the maturation of Atg8s to enable conjugation to PE, but can also reverse the reaction via de-lipidation (Agrotis et al., 2019; Kauffman et al., 2018). Atg8s were initially thought to be essential for autophagosome formation. However, cells lacking Atg8s can form small autophagosomes that fail to undergo lysosomal fusion (Nguyen et al., 2016; Vaites et al., 2018). Given that Atg4 function during autophagy is thought to be strictly mediated via processing Atg8s, it was expected that the loss of Atg4s would mimic the loss of Atg8s. In contrast, we have shown an unexpected role for the Atg4 family in phagophore membrane growth during the lipid transfer phase by promoting phagophore-ER membrane contacts (Figure 6). This discovery was made possible through the development of a new 3D reconstruction approach that we have termed AIVE which utilizes AI and the full dynamic range of EM data. It is notable that the Atg8 independent function of Atg4s during phagophore growth does not require their proteolytic activity. Given that Atg2B and Atg9 function together to drive phagophore-ER contacts (Gomez-Sanchez et al., 2018), it is possible that Atg4s may directly or indirectly influence the activity of these factors.

Atg4 protein proximity networks point to redundant and non-redundant roles for individual Atg4 family members during Atg8 independent phagophore growth. These networks include membrane and protein trafficking including Atg9 vesicles, and omegasome formation. The Atg4 proximity networks also indicate roles for Atg4s beyond autophagy, opening the door to a wider view of Atg4 activity in cell biology. There is a growing appreciation for non-autophagic roles of autophagy proteins including Atg8 conjugation factors (Galluzzi and Green, 2019). One example of a non-autophagic and Atg8 independent role for the Atg5-Atg12/Atg16L1 conjugation complex includes promoting the antiviral activity of interferon gamma (Hwang et al., 2012). Whether Atg4s can also function in cellular processes independent of their proteolytic activity remains to be determined.

Expression of an Atg8 mutant that bypasses the need for Atg4 processing (GABARAPL1-G) restored mitophagy in cells lacking Atg4s, but at a much-reduced efficiency (Figure 2C). Thus, although the Atg8 independent role of Atg4s is important for phagophore growth, it is not essential. This finding leads to a new model of autophagosome formation by the Atg8 system (Figure S9), whereby Atg4s and Atg8s play complementary yet independent roles in growing phagophore membranes. Atg4s play dual roles in this new model; 1) they promote phagophore growth by proteolytically processing Atg8s to allow attachment of Atg8s to PE, and 2) they directly promote phagophore growth independently of the proteolytic activity by driving phagophore-ER contacts. Deletion of Atg4s results in the loss of both modes of phagophore growth whereas loss of Atg8s results in the loss of only one mode. The model explains why autophagosomes can form in the absence of Atg8s but not in the absence of Atg4s.

*In vitro* studies have convincingly shown that human Atg4s can de-lipidate Atg8s (Kauffman et al., 2018), but in cells, Atg4 de-lipidation was dispensable for Atg8 recycling during autophagosome maturation (Figures 2D and S3A). Atg4 proteolytic activity is inhibited at growing phagophores by the ULK1 kinase complex which excludes a role for de-lipidation during this stage (Pengo et al., 2017; Sanchez-Wandelmer et al., 2017). It is likely that switching off de-lipidation activity in the proximity of phagophores enables Atg4s to function non-proteolytically during phagophore growth. The primary purpose of Atg4 de-lipidation activity may therefore be to prevent the basal accumulation of Atg8s on cellular membranes, including the Golgi (Figure S2H)(Joachim et al., 2015). Another important Atg8 de-conjugating activity of Atg4s lies in their ability to regulate the levels of Atg8ylation on cellular proteins including Atg16L1. Atg8ylation is a newly identified post-translational modification with the potential to play critical roles in regulating numerous cellular processes including autophagy. The future of this area of research raises new prospects into the plethora of LIR mediated complexes that can form with Atg8ylated species and how they might regulate protein function.

## Supplemental Information and Figures

### Materials and Methods

#### Cell culture, antibodies and reagents

All cell lines in this study were cultured in DMEM supplemented with 10 % (v/v) FBS (Cell Sera Australia), 1 % Penicilin-Streptomycin, 25 mM HEPES, GlutaMAX (Life Technologies) and non-essential amino acids (Life Technologies). The transfection reagents including Lipofectamine LTX (Life Technologies) and X-tremeGENE 9 (Roche) were used according to manufacturers’ instructions. The following rabbit monoclonal and polyclonal antibodies were used: Atg4C (Abcam), GABARAPL1 (Abcam), GABARAPL2 (Abcam), WIPI2 (Abcam), Atg4A (Cell Signaling), Atg4B (Cell Signaling), Atg13 (Cell Signaling), Atg16L1 (Cell Signaling), GABARAP (Cell Signaling), GABARAPL1 (Cell Signaling), GM130 (Cell Signalling), LC3A (Cell Signaling), LC3B (Cell Signaling), LC3C (Cell Signaling), HA (Cell Signaling), LAMP1 (Cell Signaling), Myc (Cell Signaling) and Tom20 (Santa Cruz). Mfn1 and B17.2L antibodies were generated previously (Lazarou et al., 2007; Lazarou et al., 2015). The polyclonal Mfn1 and B17.2L antibodies were generated in rabbits using recombinant GST-Mfn1 (aa 667-741) and full length B17.2L immunogens respectively. The following mouse monoclonal antibodies were used: CoxII (Abcam), Hsp60 (Abcam), Actin (Cell Signaling), p62 (Abnova), Parkin (Santa Cruz), Tom20 (Santa Cruz). Chicken anti-GFP and rat anti-mCherry (ThermoFisher) were also used. See Table S3 for catalogue numbers.

#### Generation of knockout lines using CRISPR/Cas9 gene editing

Atg8 hexa KO cells were described previously(Nguyen et al., 2016). Atg4 tetra KO and Atg4/Atg8 deca KO cells were generated using CRISPR guide RNAs (gRNAs) that target a common exon of all splicing variants of each Atg4 gene. Oligonucleotides (Sigma) that contain CRISPR sequences were annealed and ligated into BbsI-linearised pSpCas9(BB)-2A-GFP vector (Ran et al., 2013) (a gift from Feng Zhang; Addgene plasmid # 48138). Sequence-verified gRNA constructs were then transfected into HeLa cells for 24 h and GFP-positive cells were individually sorted by fluorescence activated cell sorting (FACS) into 96 well plates. Single cell colonies were screened for the loss of the targeted gene product by immunoblotting. The presence of frameshift indels in the genes of interest in KO clones from immunoblotting was confirmed by Sanger sequencing. Genomic DNA was first isolated and PCR was performed to amplify the targeted regions that were subsequently cloned into a pGEM4Z vector for sequencing analysis (see Table S2 for details of genotyping primers). Where antibodies were not available, three primer PCR approach(Yu et al., 2014) was employed to identify putative KO clones and then sequencing analysis was used to determine the presence of frameshift indels. Multiple knockout lines were generated by sequential transfections of one or multiple gRNA constructs. For generation of Atg4 tetra KO, Atg4D KO was first made by transfecting Atg4D CRISPR into WT HeLa cells. Atg4B and Atg4C CRISPRs were then simultaneously introduced into Atg4D KO, generating Atg4B/Atg4C/Atg4D TKO. The introduction of Atg4A CRISPR into Atg4B/Atg4C/Atg4D TKO produced Atg4 tetra KO. For Atg4/Atg8 deca KO, Atg8 hexa KO was first transfected with Atg4B CRISPR followed by Atg4A and then Atg4C CRISPRs. Atg4D CRISPR was introduced last into Atg4A/Atg4B/Atg4C/Atg8 nona KO to create Atg4/Atg8 deca KO.

#### Cloning and generation of stable cell lines

Universal vector backbones pBMN mEGFP-C1, pBMN mCherry-C1, pBMN iRFP-C1 and pBMN HA-C1 were described previously(Padman et al., 2019). Open reading frames (ORFs) of Atg4A, Atg4B, Atg4C, Atg4D and Atg2B were amplified from cDNA library with gene-specific primers and subsequently introduced into pBMN using Gibson Cloning kit (New England Biolabs). HeLa cDNA library was generated by cDNA synthesis using Random Hexamer primers (SuperScript III First-Strand Synthesis System^TM^, Thermo Fisher) and TRIzol^TM^ (Invitrogen) isolated total RNA as per manufacturers’ specifications. Atg4A-D were also individually cloned into a modified pcDNA/FRT vector harbouring a Myc-APEX2 tag by Gateway cloning (ThermoFisher Scientific). pBMN-Myc-APEX2-Atg4B was also generated.

The following plasmids were generated by ligating ORFs amplified by PCR into linearized pBMNZ using the Gibson Cloning kit: -HA-LC3A-G, -HA-LC3B-G, -HA-LC3C-G, -HA-GABARAP-G, -HA-GABARAPL1-G, -HA-GABARAPL2-G. All constructs were sequence-verified. pBMN-YFP-Parkin was described previously (Lazarou et al., 2015). Stably transfected cell lines were generated using retroviral systems as described previously(Lazarou et al., 2015), and protein expression levels were matched amongst cell lines by FACS.

#### Translocation and mitophagy and starvation treatments

For translocation and mitophagy experiments, cells were either left untreated or treated with 10 μM Oligomycin (Calbiochem), 4 μM Antimycin A (Sigma) and 5 μM QVD (ApexBio) in full growth medium for different time points as indicated in figure legends. Long treatment time point samples were treated with 10 μM QVD instead of 5 μM. For starvation experiments, cells were fed in full medium for 1 h prior to 8 h starvation using Earle’s Balanced Salt Solution (EBSS; Life Technologies).

#### Immunoblotting

HeLa cells were seeded in 6-well plates for 24 h prior to incubation with either fresh growth medium, EBSS starvation medium, or fresh medium containing 10 μM Oligomycin (Calbiochem), 4 μM Antimycin A (Sigma) and 10 μM QVD (ApexBio) for indicated times. Cells were lysed in 1× LDS sample buffer (Life Technologies) in the presence of 100 mM dithiothreitol (DTT; Sigma) and heated at 99 °C with shaking for 7 min. Approximately 25–80 μg of protein per sample was subjected to 4–12% Bis-Tris gels (Life Technologies) according to manufacturer’s instructions and electro-transferred to polyvinyl difluoride membranes (PVDF) prior to immunoblotting using indicated antibodies.

#### USP2 treatment and denaturing co-immunoprecipitation

For USP2 treatment, cells fully fed with growth medium prior to harvesting were lysed in lysis buffer (1% TX-100, 50 mM Tris-Cl pH 7.5, 150 mM NaCl, 5 mM DTT and 1 mM EDTA) on ice for 15 min. Cell lysates were then diluted with the lysis buffer without TX-100 so that the final concentration of TX-100 was 0.2 %. Each cell lysate was equally divided into two Eppendorf tubes; one of which was left untreated whereas the other was supplemented with 1 µl of recombinant USP2 (BostonBiochem). After that, all samples were incubated at 37 °C for 1 h, subjected to TCA precipitation and immunoblotting. For denaturing co-immuoprecipitation, cells were lysed in lysis buffer (1 % SDS, 50 mM Tris-Cl pH 7.5, 150 mM NaCl) supplemented with cOmplete cocktail inhibitor (Roche Applied Science) and heated at 99 °C for 5 min. Cell lysates were cleared by centrifugation at 20,000x g for 10 min at 4 °C. The collected supernatants were further diluted in 0.2 % TX-100, 50 mM Tris-Cl pH 7.5, 150 mM NaCl to make the final concentration of SDS at 0.1 % and incubated with anti-HA magnetic beads (ThermoFisher). Following 2 h incubation at 4 °C, the beads were washed three times with wash buffer (0.1% TX-100, 50 mM Tris-Cl pH 7.5, 150 mM NaCl) before eluted with 1× LDS sample buffer (Life Technologies) and analysed by immunoblotting.

#### Proximity labeling and mass spectrometry

Proximity labeling was performed as described before (Le Guerroue et al., 2017). Briefly, cells were incubated with 500 µM Biotin-Phenol (ApexBio) at 37 °C for 30 min and subsequently pulsed by addition of H2O2 for 1 min at room temperature. Afterwards, they were washed 3 times with quencher solution (10 mM sodium azide, 10 mM sodium ascorbate, 5 mM Trolox in PBS) to stop the biotinylation reaction and 3 times with PBS. Further steps were conducted at 4 °C unless stated otherwise. Cells were scraped, harvested and spun down. The resulting cell pellets were lysed in RIPA (50 mM Tris, 150 mM NaCl, 0.1 % SDS, 1 % Triton X-100, 0.5 % sodium deoxycholate) supplemented with 10 mM sodium ascorbate, 1 mM sodium azide, 1 mM Trolox and protease inhibitors (Roche Complete). Samples were sonicated 2x for 1 s, spun down at 10,000x g for 10 min before application to streptavidin agarose resin (Thermo Scientific) and incubation with overhead shaking overnight. Streptavidin-pulldown for mass spectrometry was performed similar as described before(Le Guerroue et al., 2017). Briefly, samples were washed 3 times in RIPA buffer and 3 times in 3 M Urea buffer (in 50 mM ABC) followed by incubation with TCEP (5 mM final) for 30 min at 55 °C with shaking. Samples were then alkylated with IAA (10 mM final) for 20 min at room temperature shaking in the dark and the reaction was quenched by addition of DTT (20 mM final). Samples were washed 2 times with 2 M Urea (in 50 mM ABC) before trypsin digestion overnight at 37 °C (20 µg/ml final). Samples were spun down at 1,000x g for 1 min and supernatants were collected. The resin was washed twice with 2 M Urea and supernatants were pooled with the first before acidification with TFA (1 % final). Digested peptides were desalted on custom-made C18 stage tips. Using an Easy-nLC1200 liquid chromatography, peptides were loaded onto 75 µm x 15 cm fused silica capillaries (New Objective) packed with C18AQ resin (Reprosil-Pur 120, 1.9 µm, Dr. Maisch HPLC). Peptide mixtures were separated using a gradient of 5 %–33 % acetonitrile in 0.5 % acetic acid over 90 min and detected on a Q Exactive HF mass spectrometer (Thermo Scientific). Dynamic exclusion was enabled for 30 s and singly charged species or species for which a charge could not be assigned were rejected. MS data was processed and analyzed using MaxQuant (v1.6.0.1) (Cox and Mann, 2008) and Perseus (v1.5.8.4) (Tyanova et al., 2016). All proximity experiments were performed in triplicates and the abundance of assembled proteins was determined using label-free quantification. Matches to common contaminants, reverse identifications and identifications based only on site-specific modifications were removed prior to further analysis. Student t-test was used to determine the statistical significance of the abundance alterations of biotinylated proteins between treatments (OA vs. untreated or OA/Wortmannin vs. untreated). A p-value ≤ 0.05 was considered statistically significant. Annotation enrichment analysis was performed using the PANTHER classification system (v. 14.1) (Mi et al., 2019). Similar gene ontology (GO) terms in the biological process subcategory were condensed into the broader GO-derived categories displayed in Figure 4D. The p-values displayed in Figure 4D represent the GO term that comprised the largest portion of each GO-derived category. GO terms included in each GO-derived category can be found in Table S5.

#### Immunofluorescence

For immunofluorescence, cells were cultured on HistoGrip (ThermoFisher) coated glass coverslips for 48 h before experimental treatment. All steps were performed at room temperature. Samples were first fixed with 4% (w/v) paraformaldehyde (PFA) in 0.1 M phosphate buffer (10 min), rinsed three times with PBS, permeabilized with 0.1% (v/v) Triton X-100 in PBS (10 min) and then blocked with 3% (v/v) goat serum in 0.1% (v/v) Triton X-100/PBS (15 min). The samples were incubated with indicated primary antibodies made up in 3 % (v/v) goat serum in 0.1 % (v/v) Triton X-100/PBS for 90 min and rinsed three times with PBS prior to 1 h incubation with secondary antibodies conjugated to AlexaFluor-488, Alexa-Fluor-555, AlexaFluor-633, or AlexaFluor-647 (ThermoFisher). After being washed once with 0.1 % (v/v) Triton X-100/PBS and three times with PBS, the coverslips were mounted using a TRIS buffered DABCO-glycerol mounting medium. For WIPI2b staining (Figure 3E), fixation was done by adding pre-warmed PFA to growth media (final concentration of PFA at 4% (w/v)). The rest of the staining protocol was performed as described above.

All samples were imaged in 3D by optical sectioning using an inverted Leica SP8 confocal laser scanning microscope equipped with an 63x/1.40NA objective (Oil immersion, HC PLAPO, CS2; Leica microsystems), with a minimum z-stack range of 1.8 µm and a maximum voxel size of 90 nm laterally (x,y) and 300 nm axially (z). All figure images were acquired at ambient room temperature using a Leica HyD Hybrid Detector (Leica Microsystems) and the Leica Application Suite X (LASX v2.0.1). All images are displayed as z-stack maximum projections.

#### Confocal image analysis

All 3D image data were processed and analysed by automated 3D image segmentation using 3D ROI manager (v3.93) and FeatureJ (v3.0.0) plugins for FIJI (v1.52q). As described previously (Padman et al., 2019), the image analysis workflow was divided into three discrete stages; object detection, object measurement and analysis. A global thresholding strategy was used to detect and segment Atg13 foci, WIPI2b foci, or GFP-Atg2B foci, to account for the lack of foci in the untreated conditions. Each experimental repeat was normalized by assembling a montage of maximum intensity projections for all images in a given experiment, and calculating new minimum and maximum intensity values required for linear histogram normalization of that montage. The minimum and maximum intensity values were defined if the number of pixels at a given intensity exceeded 0.02 % of all pixels in the montage. The histogram of each individual image was then rescaled between these new maximum and minimum values. The normalized data then processed with a 3D noise filter (3D median filter; 1.5 voxels) before extracting the 3D ROIs using a global threshold (64 for Atg13 and GFP-Atg2B and 96 for WIPI2b, arbitrary) via simple 3D thresholding (300 voxel maximum size). Each individual 3D ROI was applied to the original corresponding image to enable measurement of volumes and fluorescence intensities for each segmented object in all available channels. All measurements and ROIs were stored for subsequent analyses. Foci volume and number for Atg13, WIPI2b and GFP-Atg2B were quantified by direct measurement of these parameters from the 3D ROIs. The number of cells per image was quantified manually.

#### Mito-Keima autophagosome-lysosome fusion mitophagy assay

Following mitophagy induction at indicated time points, cells were trypsinised and resuspended in sorting buffer (10 % v/v FBS and 0.5 mM EDTA in PBS). After that, samples were analysed using the FAVSDiva software on a LSR Fortessa X-20 cell sorter (BD Biosciences). Lysosomal mtKeima was measured using dual excitation ratiometric pH measurements at 488 (pH 7) and 561 (pH 4) nm lasers with 695 nm and 670 nm detector filters respectively. Additional channels used include GFP (Ex/Em; 488 nm/530 nm) or iRFP670 (Ex/Em; 628 nm/670 nm). For each sample, 20,000 events were collected. Data were analysed using FlowJo (version 10). For each tested cell line, a control mtKeima ratio gate was drawn around untreated samples and was used to determine lysosomal mtKeima in treated samples; cell populations vertically shifted above the gate were classified as undergoing autophagosome-lysosome fusion and mitophagy.

#### Transmission Electron Microscopy (TEM) imaging and quantification

Samples were fixed with prewarmed phosphate-buffered 4 % PFA at 37 °C for 1 h before overnight post-fixation with 2.5 % glutaraldehyde in 0.1 M sodium cacodylate buffer at 4 °C. After 3 rinses with 0.1 M sodium cacodylate buffer; all subsequent stages were microwave assisted using a BioWave Pro microwave system (Pelco). The samples were osmicated (1 % (w/v) OsO4, 1.5 % (w/v) K3Fe(CN)6 in 0.1 M cacodylate buffer (pH 7.4)) for 1 hour at 4 °C, before exposure to three microwave duty-cycles (120 s on, 120 s off) at 100 W under vacuum. The samples were rinsed twice with MilliQ water and *en bloc* stained with 2 % (w/v) aqueous uranyl acetate with three 100 W microwave duty-cycles (120 s on, 120 s off) under vacuum. After two rinses with MilliQ water the cells were scraped, resuspended in 500 µL MilliQ, and pelleted (10,000x g). The supernatant was aspirated before resuspending and repelleting the cells in 300 µL low-melting point agarose at 37 °C. The solidified agarose-cell pellet was divided into 1 mm cubes using a razor blade for subsequent processing. Microwave assisted dehydration was performed by graduated ethanol series (80 %, 90 %, 95 %, 100 %, 100 % (v/v); each at 150 W for 40 s) and propylene oxide (100 %, 100 % (v/v); each at 150 W for 40 s). The samples were infiltrated with Araldite 502/Embed 812 by graduated concentration series in propylene oxide (25 %, 50 % 75 % 100 %, 100 % (v/v); 180 s at 250 W under vacuum), then polymerized at 60 °C. Embedded samples were sectioned using an Ultracut UCT ultramicrotome (Leica Biosystems) equipped with a 45 °C diamond knife (Diatome) to cut 75 nm ultrathin sections. The grids were stained at room temperature using 2 % (w/v) aqueous uranyl acetate (5 min) and Reynolds lead citrate (3 min) before routine imaging on a JEM-1400PLUS TEM (JEOL). For quantification, stained TEM grids were renamed and re-organised to mask their identities from the microscopist and image analyst. One ultrathin section per sample was surveyed and 15 cells were randomly selected at a low magnification (2,000x) for further analysis. Cells were excluded from selection if they were directly adjacent to a previously selected cell, partially obstructed by a grid bar, or lacking a visible nucleus. Valid cells were imaged by acquiring a montage of high magnification images, to enable the cataloguing of all visible autophagosomal structures within the cell. Data was acquired from 3 independent experiments.

#### Focussed Ion Beam-Correlative Light Electron Microscopy (FIB-CLEM)

Cells expressing MitoDsRed and GFP-Atg2B were cultured for 48 h in 35 mm Ibidi 500-grid plastic-bottomed µ-Dishes (Ibidi, Germany) prior to incubation with OA (3 h). The samples were primary fixed with prewarmed phosphate-buffered 4 % PFA at 37 °C for 1 hour, then stained (30 min) with CellMask Deep Red (ThermoFisher, 2.5 mg/ml) and Hoechst 33342 (Merck, 1 µg/ml) and rinsed with PBS prior to imaging. The fixed samples were imaged using the HyD Hybrid Detector (Leica Biosystems) of an inverted Leica SP8 confocal laser scanning microscope equipped with a 40×/1.10 objective (water immersion, HC PLAPO, CS2; Leica microsystems). The optical data (35 nm lateral pixel resolution; 200 nm axial voxel resolution) was deconvolved for subsequent alignment (fast classic maximum likelihood estimation; 10 signal-to-noise ratio; 40 iterations; 0.05 quality threshold) using Huygens Professional (v19; Scientific Volume Imaging). Navigational data was acquired by imaging three additional locations with distinct features; the motorized stage positions at these locations and the target region were extracted from the image metadata, for calculation of a linear transformation matrix defining the location of each target during later stages. After optical image acquisition, the samples were postfixed overnight with 2.5 % glutaraldehyde in 0.1 M sodium cacodylate buffer at 4 °C, then rinsed twice with 0.1 M sodium cacodylate in preparation for tertiary fixation and contrasting. All subsequent stages were microwave assisted using a BioWave Pro microwave system (Pelco). Tertiary fixation began with a modified OTO method(Seligman et al., 1966); 2 % (w/v) OsO4, 1.5 % (w/v) K3Fe(CN)6 in 0.1 M cacodylate buffer (pH 7.4); three MilliQ water rinses; 1 % (w/v) Thiocarbohydrazide in water; three MilliQ water rinses; and, 2 % OsO4 in water. Each stage used three microwave duty-cycles (120 s on, 120 s off) at 100 W under vacuum. The samples were *en bloc* stained, first with 2 % (w/v) aqueous uranyl acetate(Silva et al., 1968), then with Walton’s Lead Aspartate(Walton, 1979), with each stage using three microwave duty-cycles (120 s on, 120 s off) at 100 W under vacuum. Microwave assisted dehydration was performed by graduated ethanol series (80 %, 90 %, 95 %, 100 %, 100 % (w/v); each at 150 W for 40 s) and propylene oxide (100 %, 100 % (w/v); each at 150 W for 40 s). The samples were infiltrated with Araldite 502/Embed 812 by graduated concentration series in propylene oxide (25 %, 50 % 75 % 100 %, 100 % (v/v); 180 s at 250 W under vacuum), then polymerized at 60 °C. The positions of each target were identified from images of the resin block acquired using a Leica M125 dissection microscope with an IC80 HD camera (Leica), by using the navigational data (described earlier) to calculate the linear transformation matrix. Each target regions was physically extracted from the embedded dish after 24 h of polymerization, mounted onto 3.2 mm diameter aluminium rods with Araldite 502/Embed 812, and polymerized for an additional 48 h at 60 °C. Excess resin was trimmed from each sample using an Ultracut UCT ultramicrotome (Leica) equipped with a glass knife, before sectioning 170 µm into the Ibidi polymer-coverslip. The position of the target cells was recorded relative to distinctive markings imparted by the trimming process, by imaging the trimmed sample with the Leica M125 dissection microscope with an IC80 HD camera (Leica). The images were imported into the MAPS (v2.5; FEI) program to enable sample navigation during FIB-SEM imaging. The samples were imaged using a cryo-Helios G4 UX FIB-SEM (FEI) operating at 3.255 nm per pixel (2 kV, 100 pA, 3 µs dwell time), with ion milling at 10 nm per slice (Gallium, 30 kV, 9 nA) using the Auto Slice And View (v4.1; FEI) software. All serial block-face data was spatially registered (Rigid Transform) using the StackReg plugin for ImageJ (v1.52q) (Schindelin et al., 2012; Thevenaz et al., 1998). Orthoslices (XZ) of the aligned FIB-SEM data were spatially aligned to the deconvolved optical data using established procedures for CLEM alignment (Padman et al., 2014; Padman and Ramm, 2014).

#### FIB-CLEM processing and analyses

All image processing, reconstructions and analyses were exclusively conducted using free software under open-source or General Public licenses (GNU). Image processing was divided into three converging parallel workflows (Figure S6); Automatic membrane detection via artificial intelligence (AI), organelle sub-classification, and data normalization for voxel extraction. Automated membrane detection was performed by AI based computer vision, using the Waikato Environment for Knowledge Analysis (WEKA v3.9)(Frank et al., 2004b), in combination with training data acquired through the 3D Trainable Weka Segmentation (TWS) plugin for ImageJ(Arganda-Carreras et al., 2017; Schindelin et al., 2012). The learning model was trained in four classes of voxel; (1) Void, representing unstructured electron-lucent regions; (2) Sol, representing mid-density granular content of the cell; (3) Matter, representing non-membrane electron-dense homogenous material; and, (4) membranes. ROIs for each category of pixel were defined in cropped sub-regions of the original data to extract their numerical feature data using the 3D TWS plugin. WEKA was used to combine numerical feature data from multiple images and balance the classes (256 discretization intervals), before training the random forest classifier model for membrane detection. The trained classifier was then applied to the raw image data using the 3D TWS plugin, to extract binary masks representing the detected membranes within the dataset. The organelle sub-classification workflow was conducted manually, using the open-source Microscopy Image Browser (MIB) (Belevich et al., 2016). Image data was inspected from all available planes to ensure correct identification of organelles, which were classified into one of the following categories using the brush tool: Nucleus, early endosomes, endo-lysosomes, Golgi, microtubules, Mitochondria, and Atg2B positive phagophores (identified from the deconvolved optical data). A merged binary mask of all classified organelles was then subtracted from the membranes detected using AI; all remaining endomembranes were classified as either ER (>10,000 voxels) or vesicles (1000-10,000 voxels).

Raw voxel intensities were normalized between experiments using Contrast Limited Adaptive Histogram Equalization (CLAHE). The 3D surfaces were generated by the marching cubes algorithm with octree binning, using the open source visualization application ParaView (v5.7). Morphometric analysis of phagophore volume, structure, and the quantification of ER within 100 nm of the phagophore were also conducted in ParaView (all quantified phagophores are displayed in Figure S8). Surface meshes exported for display were decimated to reduce mesh complexity by 50 %. All surfaces were rendered using the open source rendering application Blender (v2.8) using the cycles engine.

## Acknowledgments

We thank Richard Youle for sharing plasmids and reagents, Meagan Mcgrath and Christina Mitchell for reagents, the Monash Flow cytometry Platform (FlowCore), the Monash Micro Imaging Platform, and Ramaciotti Centre for Cryo-Electron Microscopy. We also thank Gediminas Gervinskas in particular for assistance with FIB-SEM operation. This work was supported by the NHMRC (GNT1106471 and GNT1160315, M.L.), ARC future fellowship (FT1601100063, M.L.), the Deutsche Forschungsgemeinschaft (DFG) within the framework of the Munich Cluster for Systems Neurology (EXC2145 SyNergy, C.B.), the Collaborative Research Center (CRC1177, C.B.), and project grant BE 4685/2-1 (C.B.).

## Author contributions

T.N.N. and M.L. conceived the projects; T.N.N, B.P., S.Z., L.U., M.S., C.B. and M.L. designed and performed experiments; T.N.N., B.P. and M.L. wrote the manuscript and all authors contributed to preparing and editing the manuscript.

## Data availability

All data are provided in the manuscript and supplement and will be deposited on Monash University’s public data storage server (Monash.figshare).

---

**Figure S1.**
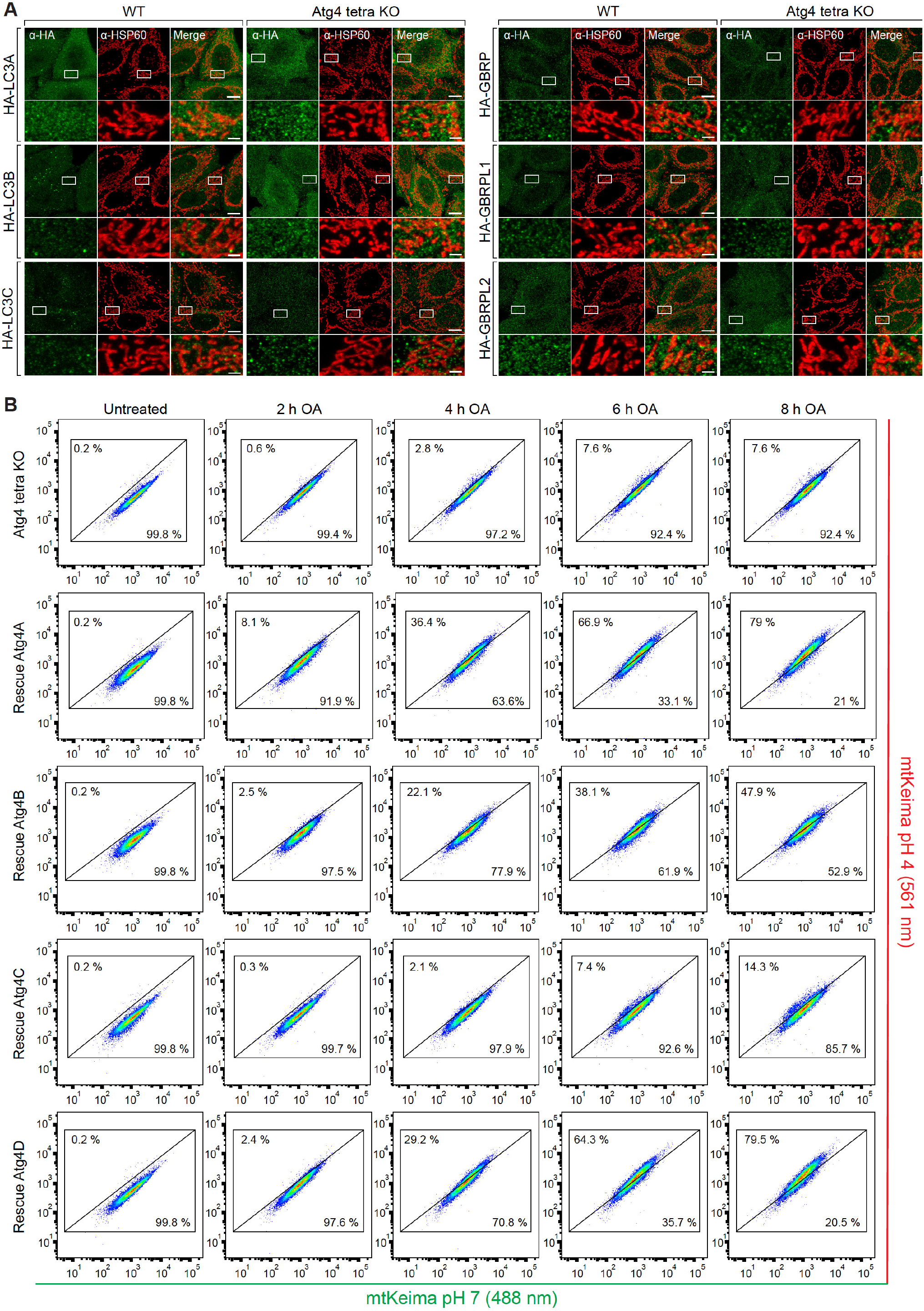
Atg4s were essential for mitochondrial recruitment of Atg8s and subsequent degradation of damaged mitochondria. (A) Representative images of untreated WT and Atg4 tetra KO cells expressing YFP–Parkin and individual HA-LC3s or HA-GBRPs immunostained for HA and HSP60. (B) Representative FACS blots for data shown in Figure 1F. Scale bars: overviews, 10 µm; insets, 2 µm.

**Figure S2.**
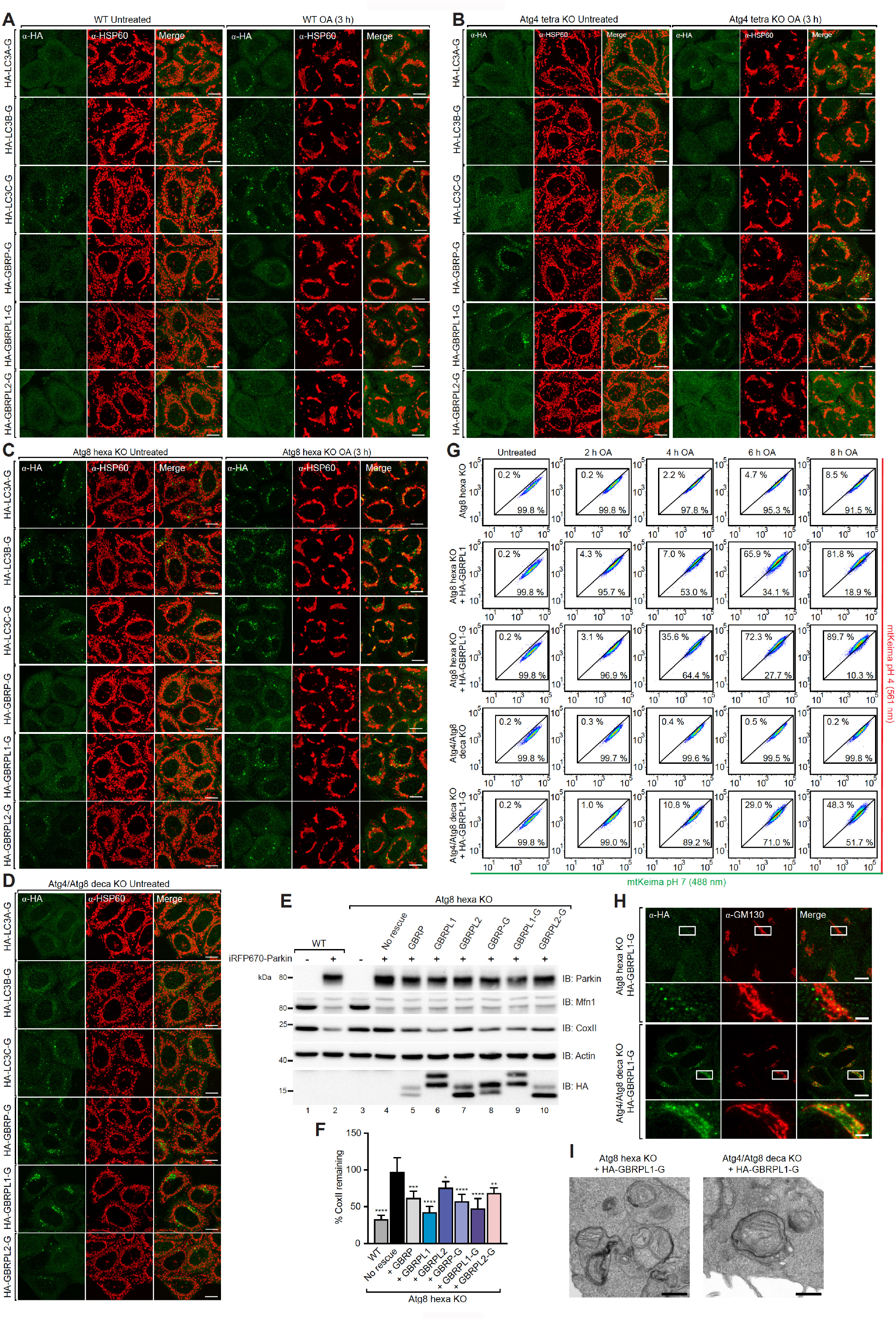
Atg8-Gs can translocate to mitochondria and drive mitochondrial degradation during PINK1/Parkin mitophagy. (A-C), Representative images of WT (A), Atg4 tetra KO (B) and Atg8 hexa KO cells (C) expressing iRFP–Parkin and individual HA-LC3-Gs or HA-GBRP-Gs untreated or treated with OA for 3 h and immunostained for HA and HSP60. (D) Representative images of untreated Atg4/Atg8 deca KO cells expressing iRFP–Parkin and individual HA-Atg8-Gs immunostained for HA and HSP60. (E and F) WT, Atg8 hexa KO cells with and without iRFP-Parkin and Atg8 hexa KO cells expressing iRFP-Parkin and individual HA-GBRPs or HA-GBRP-Gs were treated with OA for 21 h and analysed by immunoblotting (IB) (E) and CoxII levels were quantified (F). (G) Representative FACS blots for data shown in Figure 2C. (H) Representative images of untreated Atg8 hexa KO and Atg4/Atg8 KO cells expressing iRFP–Parkin and HA-GBRPL1-G immunostained for HA and GM130. (I) Representative TEM images of Atg8 hexa KO and Atg4/Atg8 deca KO cells expressing iRFP-Parkin and HA-GBRPL1-G treated with OA and BafA1 for 6 h. Scale bars: optical overviews, 10 µm; insets, 2 µm; electron micrographs, 500 nm.

**Figure S3.**
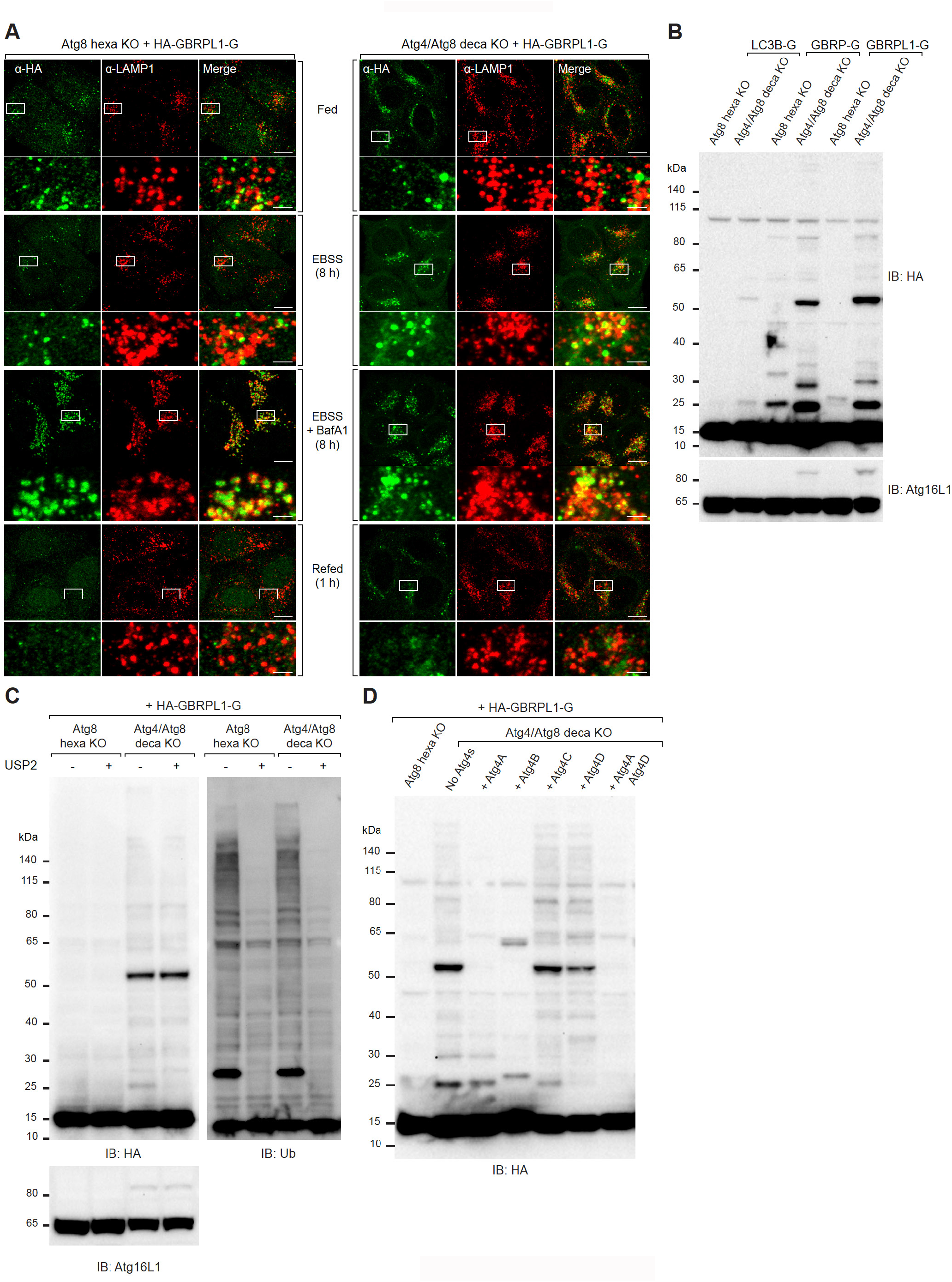
Atg4s are not required for the removal of Atg8s-PE from autolysomes but essential to prevent Atg8ylation. (A) Atg8 hexa KO and Atg4/Atg8 deca KO cells expressing iRFP-Parkin and HA-GBRPL1-G were grown in full medium or starved with EBSS in the absence or presence of BafA1 for 8 h. Following incubation, cells in EBSS without BafA1 were refed with full medium for 1 h. All samples were subjected to immunostaining with anti-HA and anti-LAMP1 antibodies. (B) Atg8 hexa KO and Atg4/Atg8 deca KO cells expressing HA-LC3B-G, HA-GBRP-G or HA-GBRPL1-G were immunoblotted for HA and Atg16L1. (C) Samples from fed Atg8 hexa KO and Atg4/Atg8 deca KO cells expressing iRFP-Parkin and HA-GBRPL1-G were incubated in the absence or presence of USP2 prior to immunoblotting analysis with indicated antibodies. (D) Atg8 hexa KO and Atg4/Atg8 deca KO cells expressing iRFP-Parkin and HA-GBRPL1-G without Atg4s or with indicated Atg4s were subjected to immunoblotting with anti-HA antibody. Scale bars: overviews, 10 µm; insets, 2 µm.

**Figure S4.**
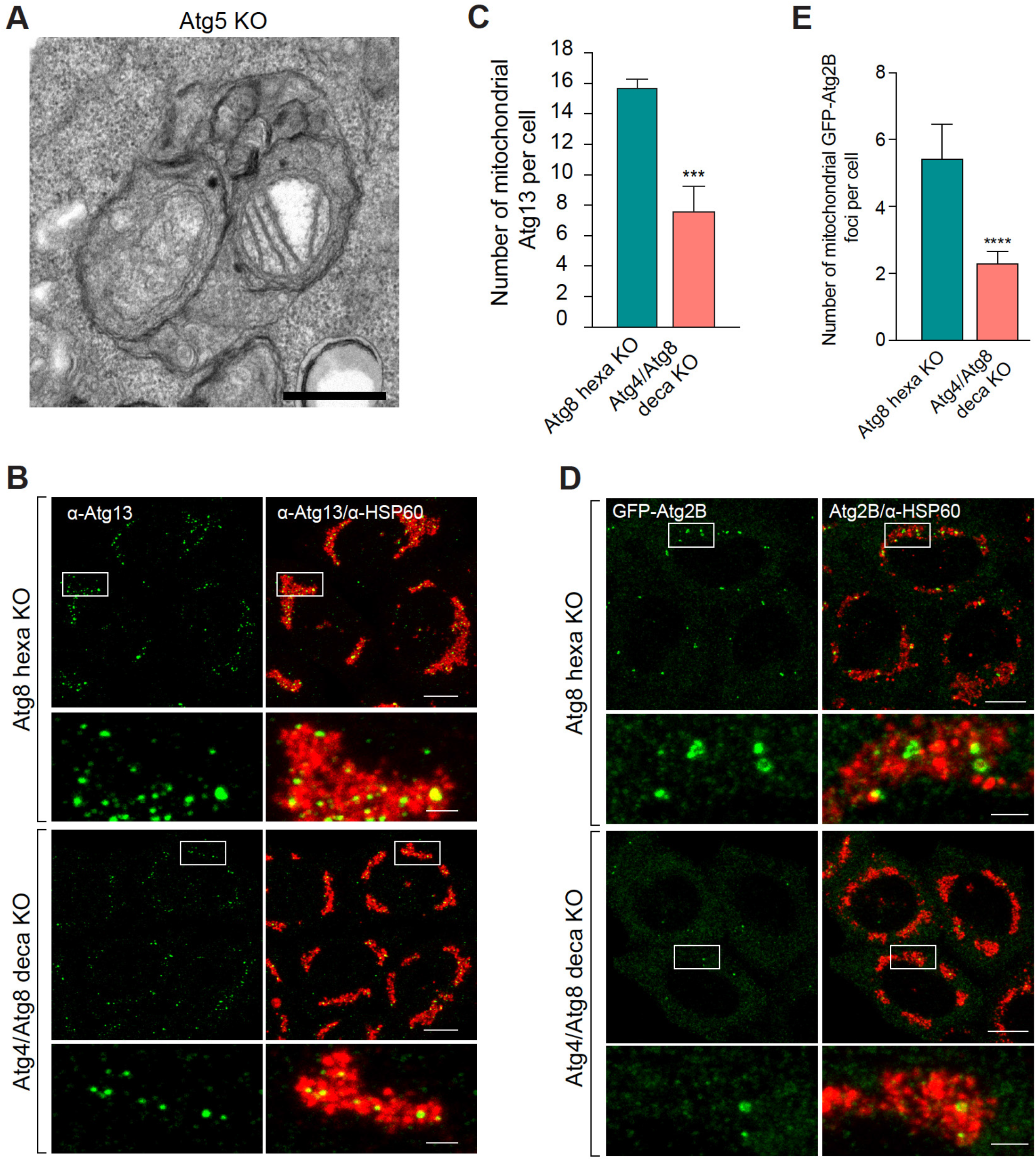
Atg4s directly drive phagophore progression independent of Atg8s. (A) Representative TEM image of Atg5 KO treated with OA and BafA1 for 6 h. (B) Representative image of Atg8 hexa KO and Atg4/Atg8 deca KO cells expressing iRFP-Parkin treated with OA for 3 h and immunostained for Atg13 and HSP60. (C) The average number of mitochondrial Atg13 structures from (C) was analysed. (D) Representative images of Atg8 hexa KO and Atg4/Atg8 deca KO cells expressing iRFP-Parkin and GFP-Atg2B immunostained for GFP and HSP60. (E) The average number of mitochondrial GFP-Atg2B structures per cell from (D) was analysed. Data in (C and E) are mean ± s.d. from three independent experiments. \*\*\*\**P*<0.0001 (one-way ANOVA). Scale bars: overviews, 10 µm; insets, 2 µm; electron micrograph, 500 nm.

**Figure S5.**
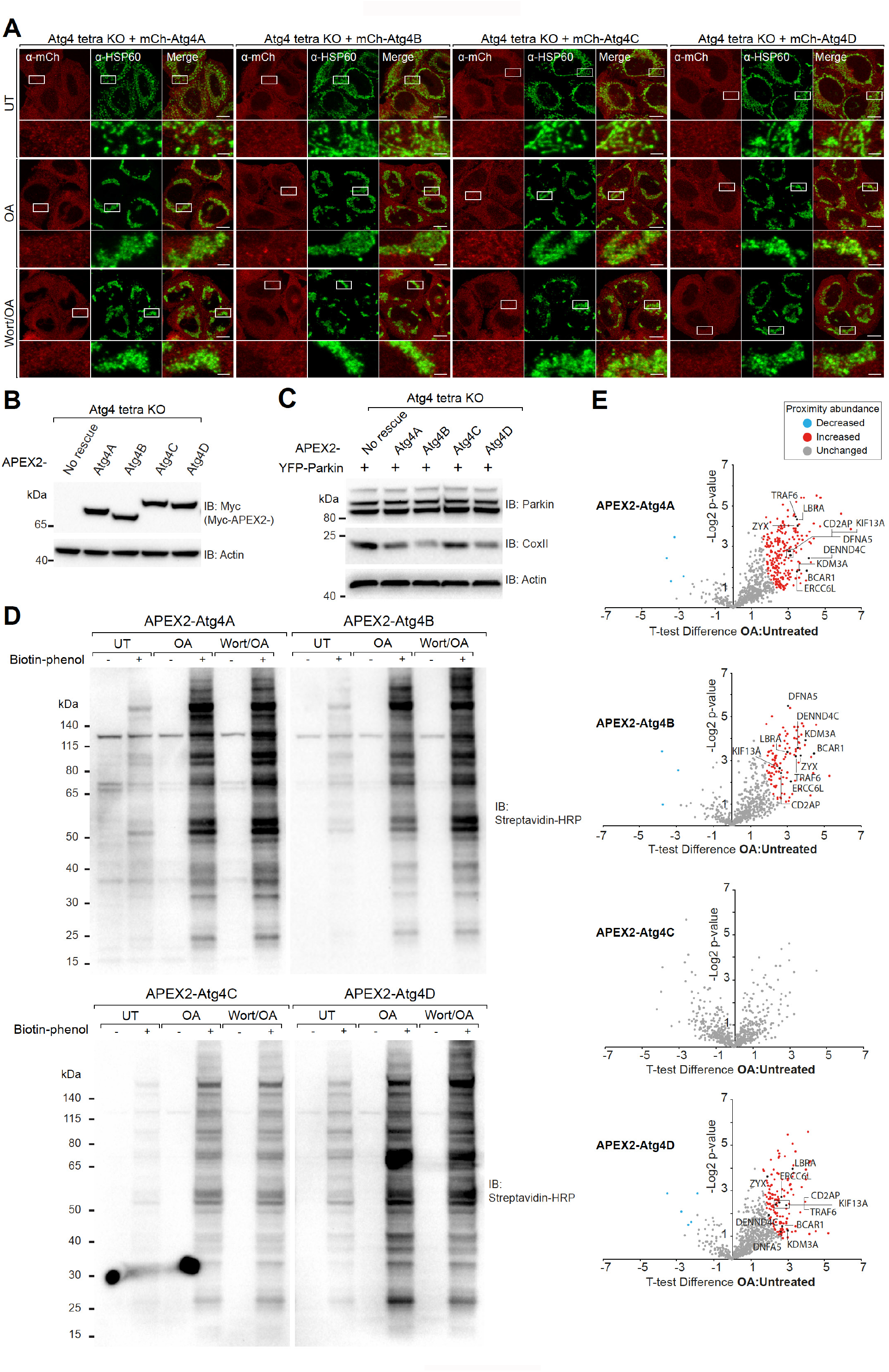
Biotinylation proximity labelling of APEX2-Atg4s. (A) Representative images of Atg4 tetra KO cells expressing YFP-Parkin and individual mCh-Atg4s left untreated, treated with OA, or OA with Wortmannin (Wort/OA) for 3 h and immunostained for mCherry and HSP60. (B) Atg4 tetra KO cells with and without re-expression of individual APEX2-Atg4s were analysed by immunoblotting. (C) Indicated cell lines were treated with OA for 21 h and immunoblotted with indicated antibodies. (D) Atg4 tetra KO cells with or without re-expression of individual APEX2-Atg4s were left untreated, treated with OA only, or Wort/OA for 3 h. Following biotinylation labelling reactions, cells were then harvested and subjected to immunoblotting with anti-Streptavidin-HRP. (E) Scatterplot analysis of enriched biotinylated targets of each APEX2-Atg4 from Figure 4C. Scale bars: overviews, 10 µm; insets, 2 µm.

**Figure S6.**
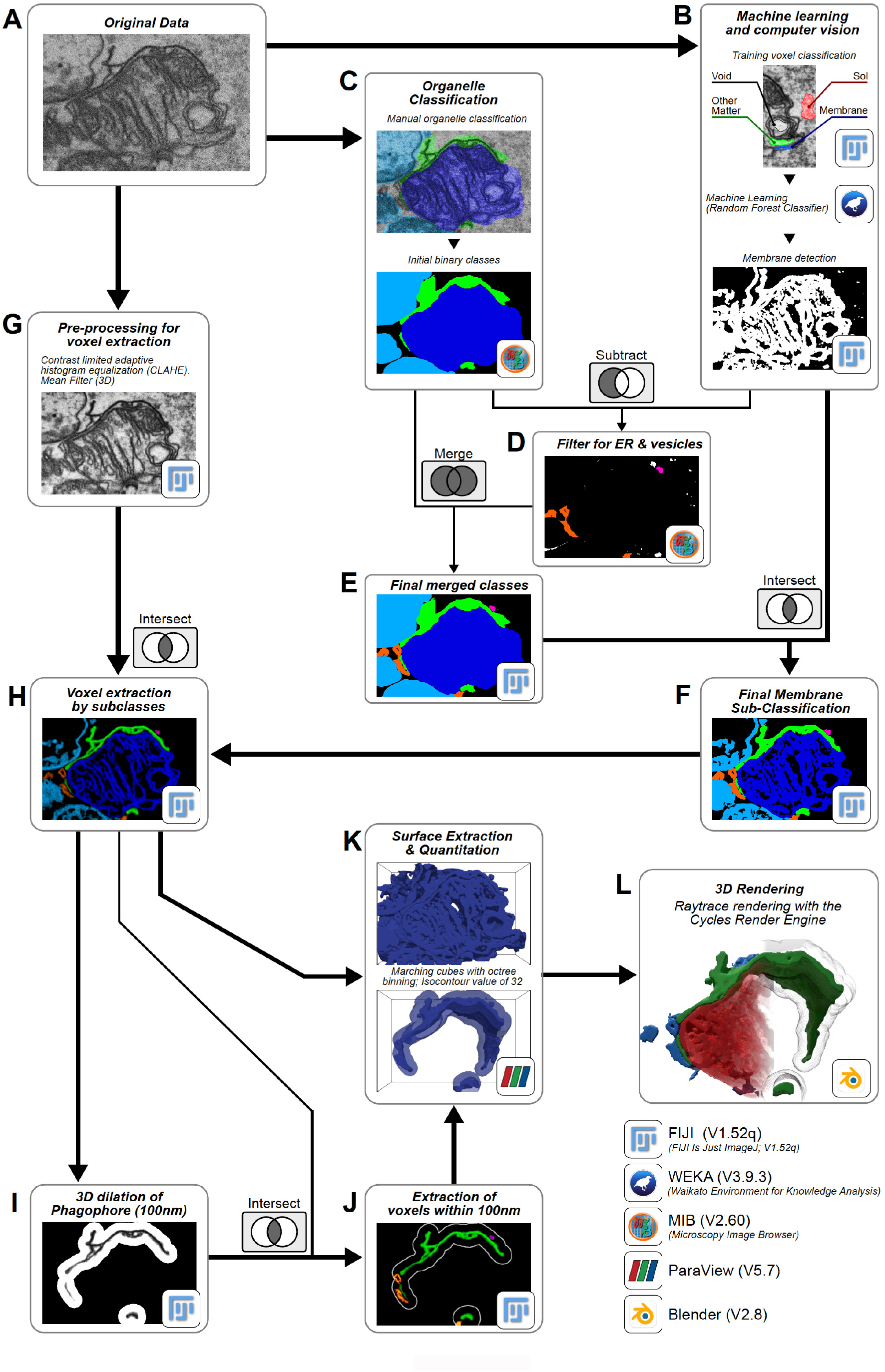
Workflow for generating FIB-CLEM based 3D reconstructions. (A) Duplicate copies of the FIB-SEM data were generated for processing via three parallel workflows: (B) machine learning and automated membrane detection using WEKA (v3.9.3) with the TWS plugin for Fiji (1.52qv), (C) manual organelle classification using MIB (v2.6), and (G) and data normalization using Fiji. (D) ER and vesicles were classified by subtracting the initial organelle classifications (C) from the detected membranes (B), and filtering all remaining endo-membranes by size and shape. (E) The ER and vesicle classes were merged with the initial organelle classifications to generate a final set of classified binaries. (F) To sub-classify the membranes belonging to each organelle, boolean intersections were calculated between the binary classes (E) and the detected membranes (B). (G) To account for potential image intensity variation between the samples, data were normalized using Contrast Limited Adaptive Histogram Normalization and mean filtered to remove detector noise. (H) Voxels representing the membranes from each sub-class of organelle were then extracted, by calculating the boolean intersect between the normalized data (G) and the sub-classified membrane binaries (F). (I and J) To quantify the volume of ER within 100 nm of the phagophore, the sub-class of voxels representing the phagophore was binarized (threshold value of 32; as used in (K)) and spherically dilated by 100 nm in 3D (I), before calculating its boolean intersect with the ER sub-class of voxels. (K) Surfaces for all organelles (H) and the ER within 100 nm of the phagophore (J) were generated in ParaView (v5.7), using the marching cubes algorithm with octree binning (isocontour value of 32); all key quantitation parameters were also extracted. (L) The surface meshes used for rendering were decimated by 50 % in ParaView (K) and imported into Blender for final 3D rendering.

**Figure S7.**
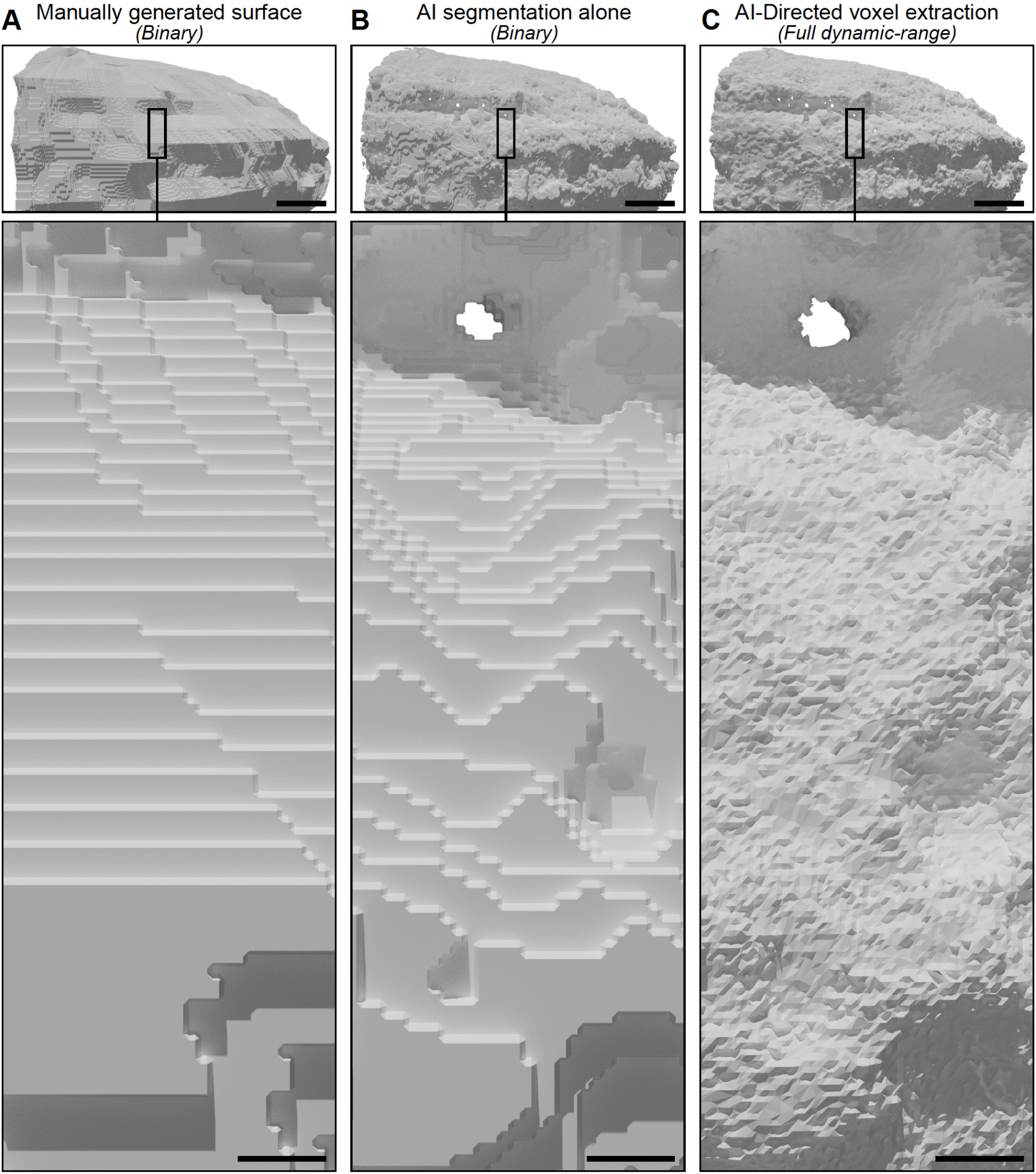
Comparison of approaches to the 3D reconstruction of FIB-SEM data. (A-C) 3D renderings of the nuclear envelope shown in Figure 5D generated via (A) traditional manual segmentation of membranes, (B) image segmentation by AI, or (C) the AI-directed approach outlined in Figure S6. Surfaces in (C) were generated by merging the data shown in A and B with the FIB-SEM data, to selectively extract the nuclear envelope voxels for reconstruction. Scale bars: overviews, 500 nm; insets, 50 nm.

**Figure S8.**
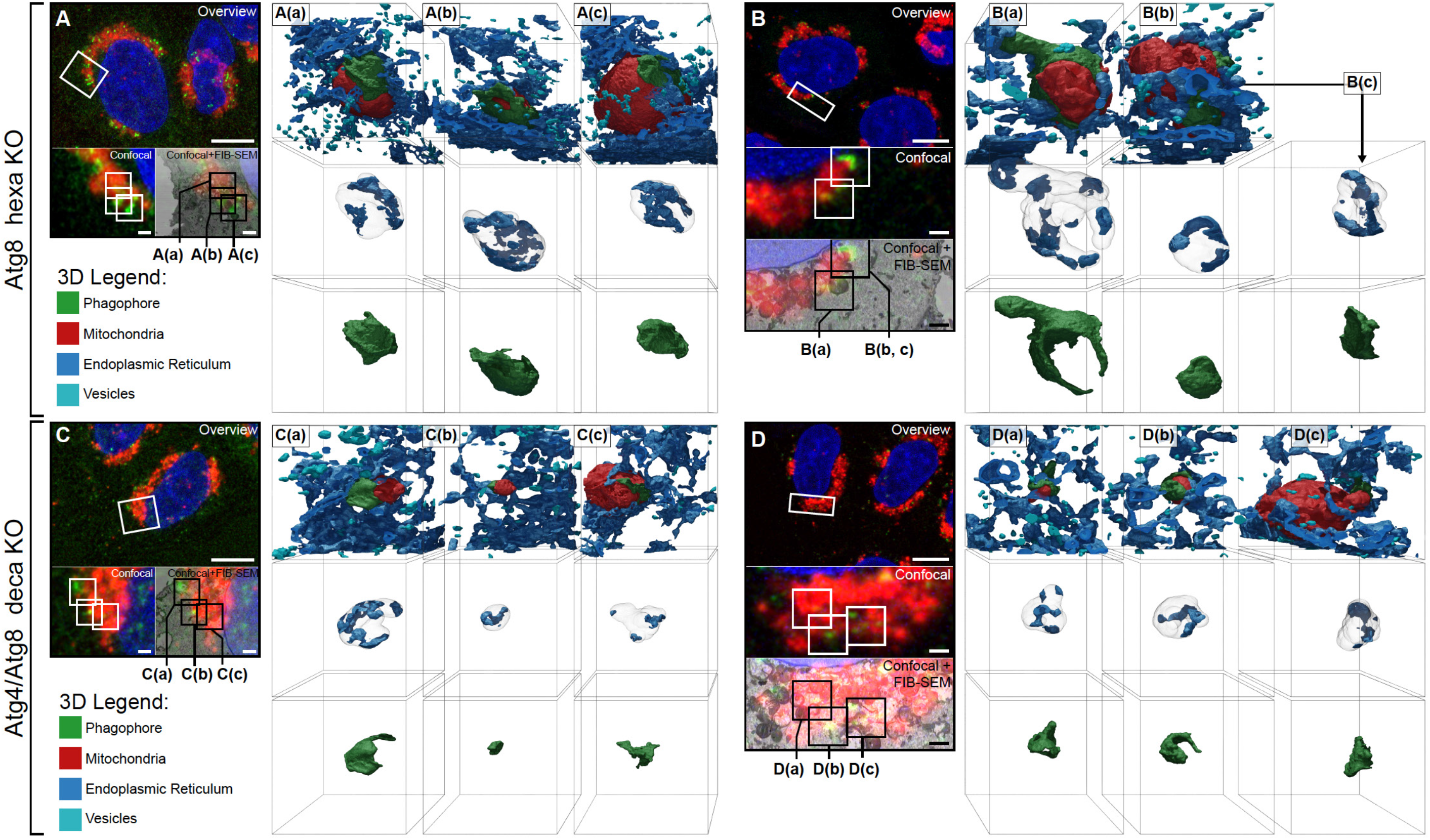
Phagophore growth and phagophore-ER contacts are significantly reduced in the absence of Atg4s. (A-D) Correlative alignment between the deconvolved optical data and orthosliced (X/Z) FIB-SEM data for all detected Atg2B positive phagophores. Indicated insets show six phagophores from two different (A and B) Atg8 hexa KO cells, or (C and D) Atg4/Atg8 deca KO cells expressing iRFP-Parkin and GFP-Atg2B after 3 h incubation with OA. Scale bars: optical overviews, 10 µm; CLEM alignment insets, 1 µm; rendered inset cubes, 2 µm x 2 µm x 2 µm.

**Figure S9.**
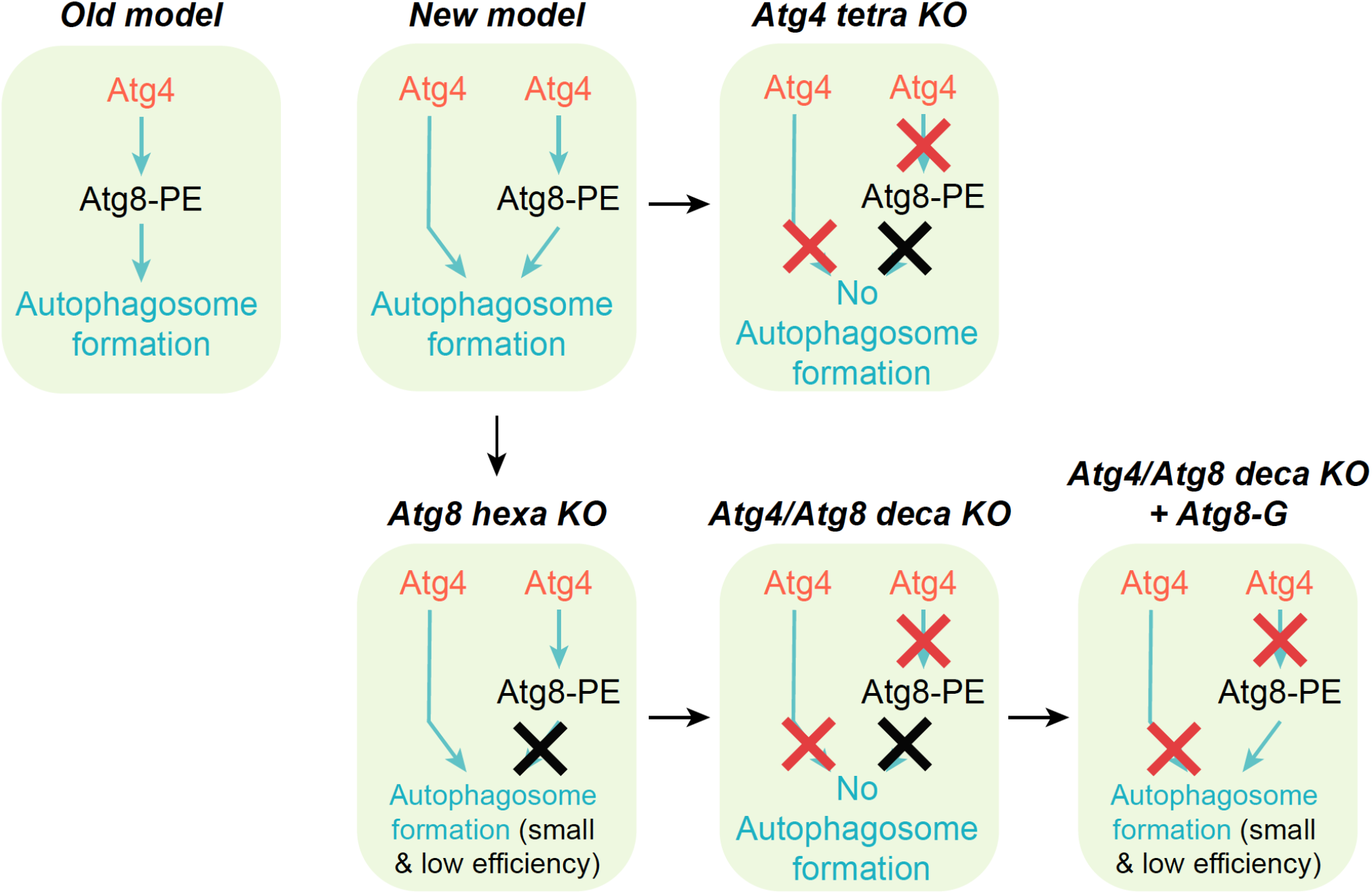
Updated model of autophagosome biogenesis. The old model dictates that autophagosome biogenesis is mediated by lipidated Atg8s or Atg8-PE. Functions of Atg4s during autophagosome formation are believed to be solely dependent on their ability to process Atg8s. In our new model, Atg4s play dual roles during autophagosome biogenesis; they can function through Atg8s (mode 1) or directly promote phagophore growth independent of Atg8s (mode 2). Therefore, autophagosomes can still be made when one of the modes is functional, i.e. in Atg8 hexa KO (functional mode 1) or in Atg4/Atg8 deca KO expressing Atg8-Gs. In the latter case, Atg8-Gs can bypass initial Atg4 process and be directly conjugated to PE so mode 2 is still functional in the absence of Atg4s. However, in both cases the generated autophagosomes are smaller and autophagosome formation efficiency is much lower compared to WT cells. When both modes are absent i.e. in Atg4 tetra KO and Atg4/Atg8 deca KO, autophagosomes can no longer be generated.

**Table S1.**
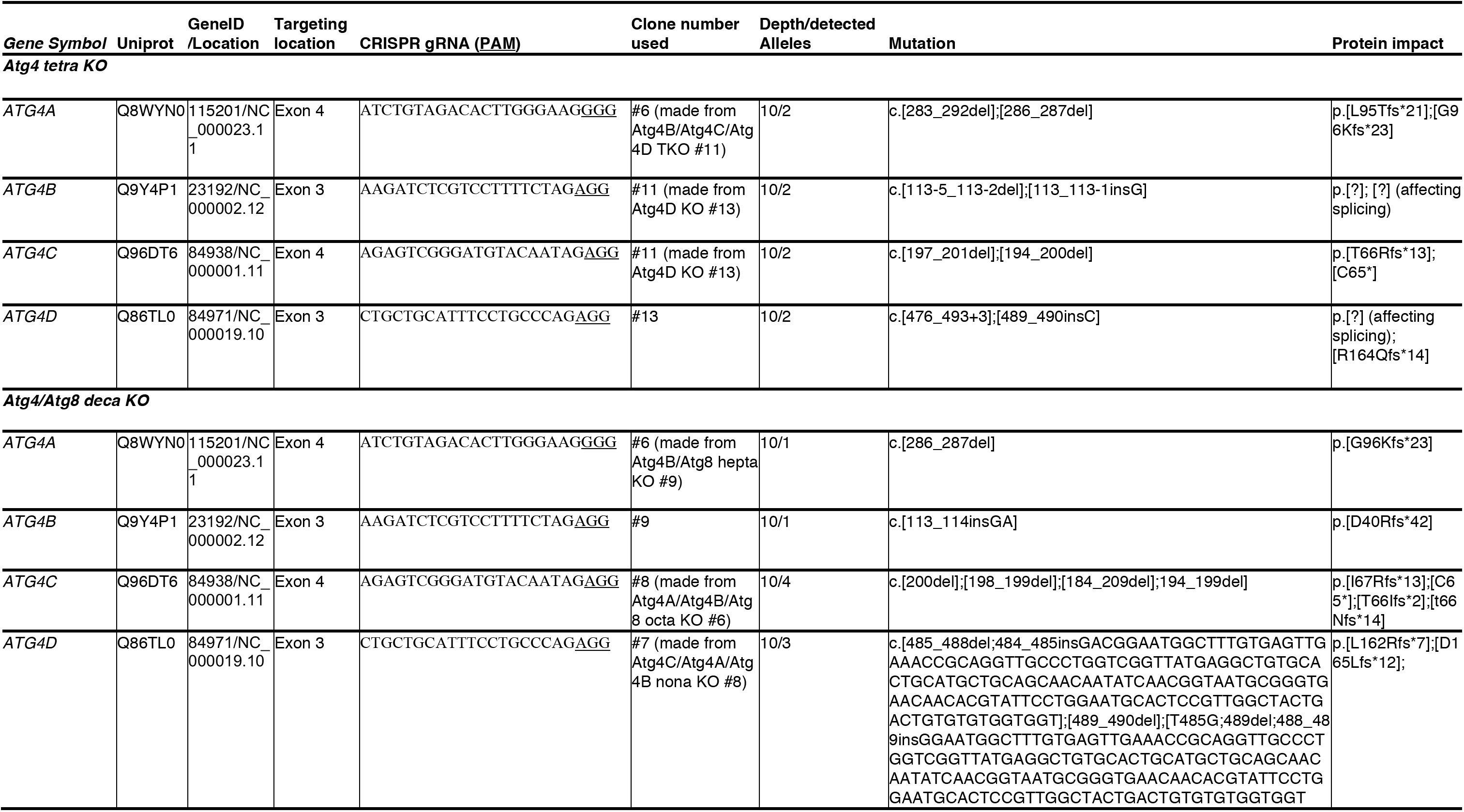
CRISPR sequences and genotyping results of Atg4 tetra KO and Atg4/Atg8 deca KO. The indels for ATG4 genes detected in the indicated knockout cell lines (“Mutation” column) and their translated proteins (“Protein impact” column) are formatted according to Human Genome Variation Society (HGVS; http://varnomen.hgvs.org/). Mutation positions within genes which have multiple splice variants are determined using the variant indicated as the canonical isoform in Uniprot. del = deletion; ins = insertion; c. = coding DNA; p. = protein; fs = frame shift; * = stop codon; [?] = affecting splicing. The numbers following the asterisks denote the numbers of amino acids between the first amino acid changed after the mutation(s) and the first subsequent stop codon encountered.

**Table S2.**
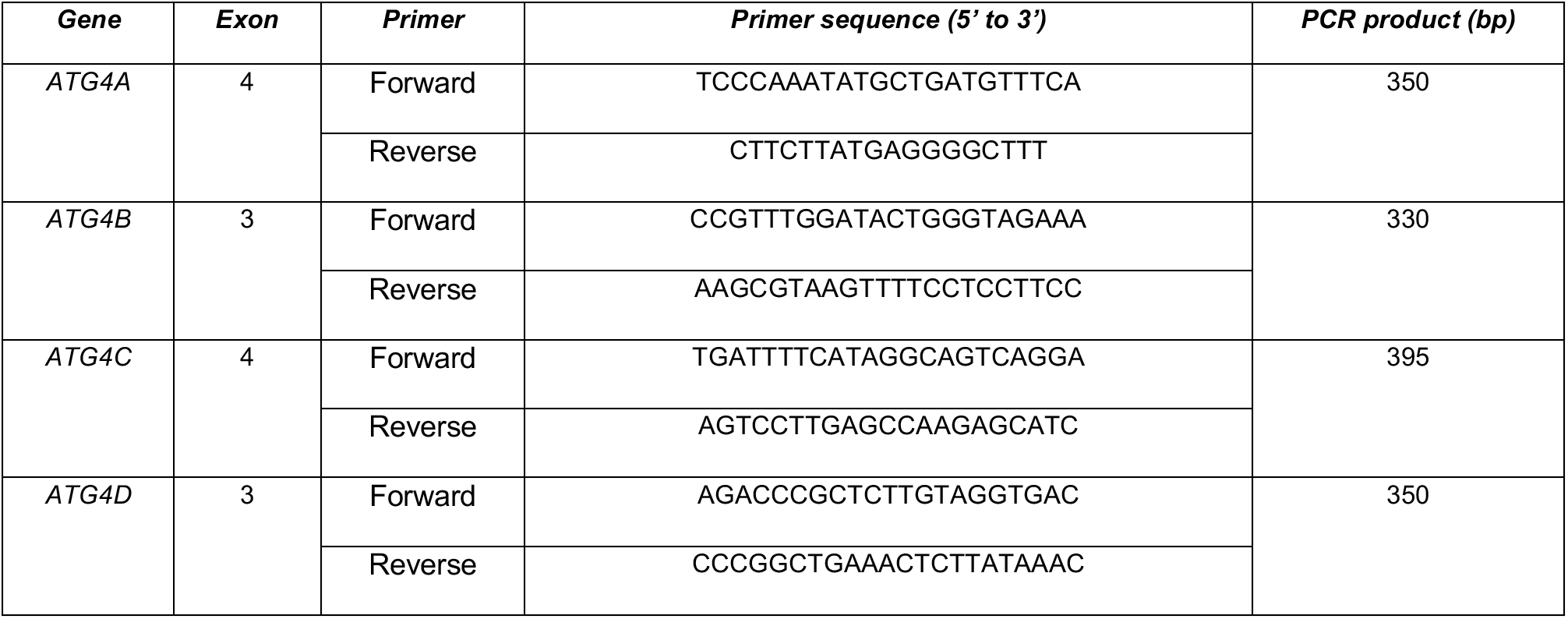
Primer used for genotyping of the generated knockout lines.

**Table S3.** Details of antibodies used in this study.

| Antibodies | Company | Species | Catalog # |
| --- | --- | --- | --- |
| Atg4C | Abcam | Rabbit | ab183516 |
| CoxII | Abcam | Mouse | ab110258 |
| GABARAPL1 | Abcam | Rabbit | ab86497 |
| GABARAPL2 | Abcam | Rabbit | ab126607 |
| WIPI2b | Abcam | Mouse | ab105459 |
| p62 | Abnova | Mouse | H00008878-M01 |
| Actin | Cell Signaling | Rabbit | 4967S |
| Atg4A | Cell Signaling | Rabbit | 7613 |
| Atg4B | Cell Signaling | Rabbit | 13507 |
| Atg13 | Cell Signaling | Rabbit | 13468 |
| Atg16L1 | Cell Signaling | Rabbit | 8089 |
| GABARAP | Cell Signaling | Rabbit | 13733S |
| GM130 | Cell Signaling | Rabbit | 12480S |
| HA | Cell Signaling | Rabbit | 3724S |
| LAMP1 | Cell Signaling | Rabbit | 9091P |
| LC3A | Cell Signaling | Rabbit | 4599S |
| LC3B | Cell Signaling | Rabbit | 4108S |
| LC3C | Cell Signaling | Rabbit | 14736S |
| Myc | Cell Signaling | Rabbit | 2278 |
| Streptavidin-HRP | Cell Signaling | NA | 3999 |
| Parkin | Santa Cruz | Mouse | sc-32282 |
| TOM20 | Santa Cruz | Rabbit | sc-11415 |
| TOM20 | Santa Cruz | Mouse | sc-17764 |
| GFP | ThermoFisher | Chicken | A10262 |
| GST | ThermoFisher | Rabbit | 71-7500 |
| mCherry | ThermoFisher | Rat | M11217 |

**Table S4. Excel table containing proteomic proteins identified from biotinylation proximity labelling of APEX2-Atg4s in triplicate.** Atg4 tetra KO cells expressing YFP-Parkin and APEX2-Atg4A (A), APEX2-Atg4B (B), APEX-Atg4C (C) or APEX2-Atg4D (D) under unstimulated (samples 1-3), OA treated (samples 4-6) or Wort/OA (samples 7-9) conditions were subjected to biotinylation labelling and MS analysis.

**Table S5. Excel table containing Raw data for Gene Ontology analysis graph (Figure 4D).**

**Movie S1.** Animated 3D rendering of Figure 5 showing all membranes detected using the computer vision procedure, with the plasma membrane removed. Larger membranes (gold) were separated from smaller vesicles (blue) by size. The nuclear envelope (purple) was defined by manual sub-classification of the corresponding membranes, and overlaid with the Cryo-EM structure of the human Nuclear Pore Complex (orange) solved *in situ* by A von Appen et al., 2015.

**Movie S2.** Animated rotation 3D rendering from Figure 6E, showing mitochondria (red) undergoing sequestration by a phagophore (green) surrounded by ER. Colours for other organelles indicated in Figure 6. Inset cube dimensions, 2 µm x 2 µm x 2 µm.

**Movie S3.** Animated rotation 3D rendering from Figure 6F, showing mitochondria (red) undergoing sequestration by a phagophore (green) surrounded by ER. Colours for other organelles indicated in Figure 6. Inset cube dimensions, 2 µm x 2 µm x 2 µm.

**Movie S4.** Animated rotation 3D rendering from Figure 6G, showing mitochondria (red) undergoing sequestration by a phagophore (green) surrounded by ER. Colours for other organelles indicated in Figure 6. Inset cube dimensions, 2 µm x 2 µm x 2 µm.

**Movie S5.** Animated rotation 3D rendering from Figure 6H, showing mitochondria (red) undergoing sequestration by a phagophore (green) surrounded by ER. Colours for other organelles indicated in Figure 6. Inset cube dimensions, 2 µm x 2 µm x 2 µm.

